# Ramping-up hippocampal ripples and their neocortical coupling support human visual short-term memory

**DOI:** 10.64898/2026.04.07.716930

**Authors:** Jing Liu, Xianhui He, Can Yang, Nikolai Axmacher, Gui Xue, Shaoming Zhang, Ying Cai

## Abstract

Emerging evidence suggests that the hippocampus contributes to visual short-term memory (VSTM). However, the neural mechanisms through which hippocampal activity supports the maintenance of VSTM representations remain largely unknown. Here, using intracranial EEG recordings from human participants performing a delayed match-to-sample task for naturalistic objects, we investigated the role of hippocampal ripple activity—brief high-frequency oscillations associated with memory replay—in VSTM maintenance and hippocampal-neocortical communication. We found that hippocampal ripple rates progressively ramped up during the maintenance period and supported successful VSTM. More critically, hippocampal ripples were temporally coupled with the ripples in the lateral temporal lobe (LTL), and these coupled ripples were associated with memory reactivation in the LTL. These findings provide direct evidence that hippocampal–neocortical interaction via coupled ripples supports VSTM, extending the role of hippocampal ripples from long-term consolidation to short-term mnemonic processes and offering new insights into the dynamic coding mechanisms of working memory.

**Impact Statement:** Hippocampal ripple ramping-up and neocortical coupling support a dynamic coding mechanism for visual short-term memory maintenance

## Introduction

Early research suggested that the hippocampus is essential exclusively for long-term episodic memory formation (Scoville and Milner, 1957; Squire and Zola-Morgan, 1991; Vargha-Khadem et al., 1997). However, emerging evidence implicates that the hippocampus also contributes to visual short-term memory (VSTM) (Li et al., 2024; Ranganath and D’Esposito, 2001; von Allmen et al., 2013). For example, patients with focal hippocampal damage show impaired VSTM (Borders et al., 2022; Koen et al., 2017; Yonelinas, 2013). Moreover, item-specific VSTM representations could be decoded or reconstructed from the hippocampal neuronal activity (Kamiński et al., 2017; Kornblith et al., 2017; Xie et al., 2023). These findings suggest that the hippocampus contributes to maintaining VSTM representations, yet the underlying mechanisms remain unclear.

A key candidate mechanism is the hippocampal ripple, a brief high-frequency (∼80-170 Hz) oscillation that reflects synchronized activity of neuronal ensembles in the hippocampus (Buzsáki, 2015; Iwata et al., 2024; Norman et al., 2019). Traditionally, hippocampal sharp-wave ripples have been recognized for their role in offline memory consolidation during sleep and quiet awake rest, during which they are temporally coordinated with neocortical memory replay/reactivation via hippocampal-neocortical communication (Eschenko et al., 2008; Schreiner et al., 2024; Zhang et al., 2018, 2024). Recent studies, however, indicate that ripples also actively support online memory processing. In human long-term memory (LTM) studies, ripple rates during encoding and pre-retrieval intervals predict subsequent LTM performance, and temporally coordinate the reinstatement of neocortical representations during successful retrieval (Kunz et al., 2024; Norman et al., 2019; Sakon and Kahana, 2022; Vaz et al., 2019; Zhang et al., 2024). Beyond LTM, rodent studies have revealed that selectively disrupting hippocampal ripples during awake learning periods impaired the working memory-dependent spatial navigation performance (Jadhav et al., 2012; Zhang et al., 2021), pointing to a vital role for ripples in sustaining online short-term memory representations. Moreover, VSTM and LTM could recruit overlapping hippocampal-neocortical networks (Cowan, 2019; Ranganath and Blumenfeld, 2005), especially when the VSTM task is demanding or engaging feature binding (Hannula et al., 2006; Jeneson et al., 2012; Jeneson and Squire, 2012; Pertzov et al., 2013). These findings together raise a fundamental question: Do hippocampal ripples, long implicated in LTM formation and offline consolidation, also actively support human VSTM maintenance?

In parallel, traditional models of VSTM proposing sustained, persistent neural activity during delay periods have been challenged by “activity-silent” or dynamic coding frameworks (Fiebig and Lansner, 2017; Miller et al., 2018; Mongillo et al., 2008; Stokes, 2015). According to these frameworks, VSTM representations are maintained via short-term synaptic plasticity that stores information in an “activity-silent” hidden state. Transient bursts are required to refresh the short-lived latent synaptic trace and thus maintain the stored representations (Mongillo et al., 2008). Moreover, these bursts are posited to “ramp up” toward the end of the retention interval, reflecting memory readout and preparation for responses (Stokes, 2015), a prediction supported by non-human primate research (Lundqvist et al., 2018b; Spaak et al., 2017). Given that hippocampal ripples share key properties postulated for bursts, such as high-frequency, discreteness, and temporal coordination with neocortical memory reactivation (Norman et al., 2019), we hypothesized that hippocampal ripple rates would exhibit a progressive temporal ramping-up pattern across the VSTM maintenance interval in accordance with the dynamic coding framework.

Furthermore, effective memory processing depends on hippocampal-neocortical communication (Gattas et al., 2023). This cross-regional communication can occur efficiently via coupled ripples, i.e., co-occurrence of hippocampal ripples and neocortical ripples (Khodagholy et al., 2017), which have been associated with successful memory retrieval (Seger et al., 2025; Vaz et al., 2019). In particular, we are interested in the interaction between the hippocampus (HPC) and the lateral temporal lobe (LTL), a region implicated in representing memory content during both episodic memory retrieval (Pacheco Estefan et al., 2019) and VSTM (Miller et al., 1993; Ranganath, 2006). We thus hypothesized that hippocampal ripples temporally couple with LTL ripples to orchestrate the reactivation of stimulus-specific representations in the LTL.

To test these hypotheses, we analyzed iEEG recordings from the HPC and LTL in neurosurgical patients performing a VSTM task that required maintaining and then discriminating target images from highly similar lures. We found that hippocampal but not LTL ripple rates ramped up during the maintenance period, supporting VSTM accuracy. Critically, hippocampal ripples were temporally coupled with LTL ripples, and these coupled ripples further coincided with memory reactivation in the LTL. Importantly, these VSTM-related ripple dynamics cannot be attributed to the formation of subsequent long-term memory. Together, these findings provide direct evidence that hippocampal ripples coordinate neocortical memory reactivation through dynamic coding mechanisms in supporting the VSTM.

## Results

### Behavioral results

Thirteen participants (mean age ± SD: 26.77 ± 5.48 years, 7 females) with drug-resistant epilepsy were included in this study. The VSTM was assessed using a delayed match-to-sample (DMS) task comprising three stages: encoding, maintenance, and retrieval (Fig. 1a, upper; see Methods). In each trial, participants encoded a word-picture pair for 3 s. This was followed by a 7-second maintenance period, during which the picture was removed, and participants were asked to vividly mentally maintain the picture. During the immediate retrieval stage, a probe picture (i.e., either the original target or a highly similar lure) was presented, and participants indicated whether it matched the encoded picture (see Fig. 1a). All pictures were from four categories (i.e., animals, fruits, electrical devices, and furniture). Participants performed well in the DMS task with an accuracy of 89.71 ± 4.26% (mean ± SD) and a response time (RT) of 1.03 ± 0.22 s (mean ± SD). In addition, the RT for correct trials (i.e., remembered trials) was significantly shorter than that for incorrect trials (i.e., forgotten trials, *t*(12) = −5.88, *p* < 0.001). Note that the cue word in the DMS task was designed for a subsequent long-term memory cued-recall test (see details in Methods).

**Figure 1.**
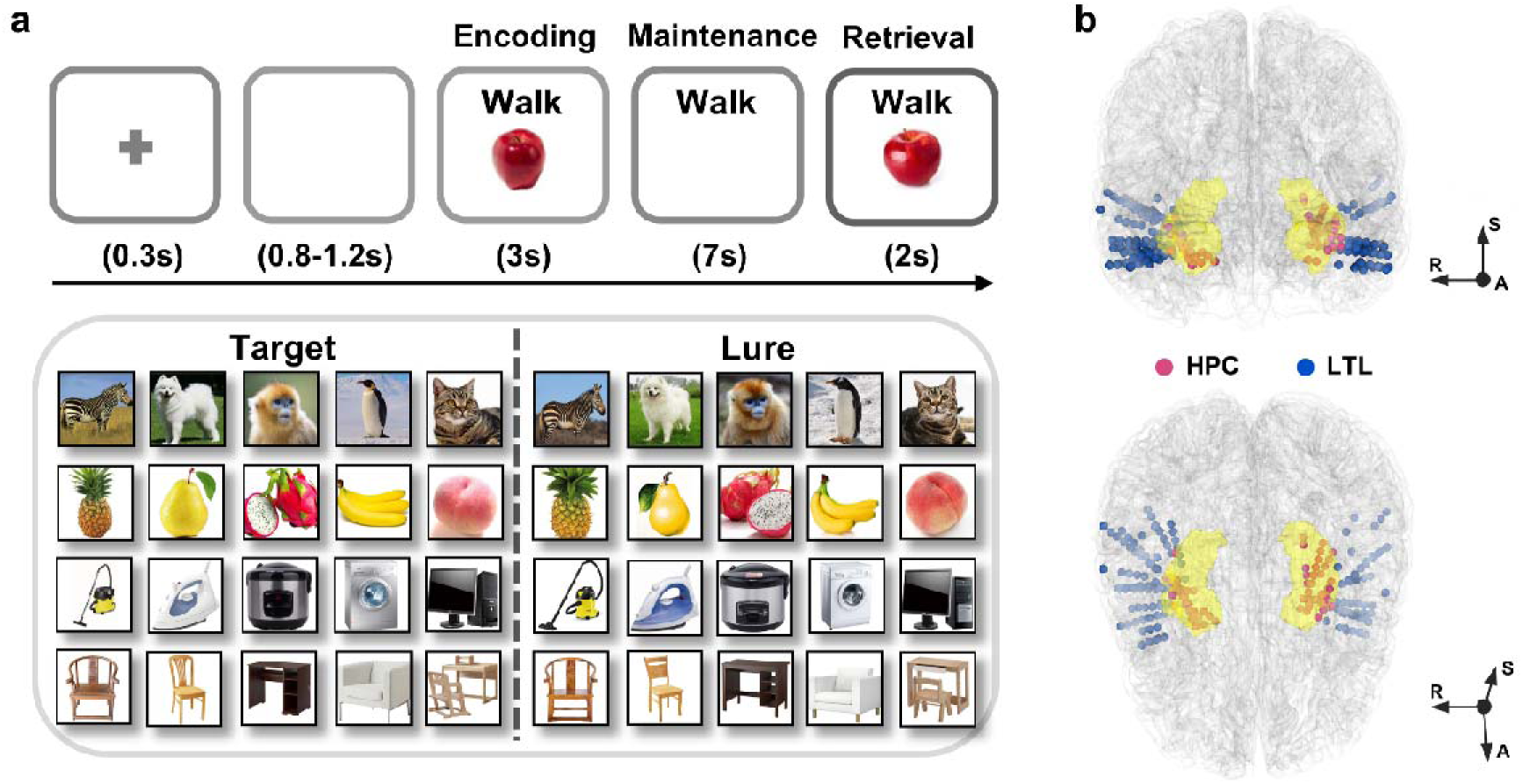
Experimental paradigm, stimuli, and intracranial EEG channel localization. **(a)** Upper: an exemplar trial procedure of the delayed match-to-sample (DMS) task. Lower: examples of target and lure pictures. Pictures were selected from four categories—animals, fruits, electrical appliances, and furniture—with each row representing one category; **(b)** Normalized locations of intracranial EEG channels across all 13 participants. Pink spheres indicate hippocampal (HPC) channels within the highlighted yellow-shaded region; blue spheres indicate lateral temporal lobe (LTL) channels.

### Hippocampal ripple rates during the VSTM task

All participants were implanted with depth electrodes for clinical purposes. The iEEG data were recorded from the hippocampus during the DMS task (69 channels, mean ± SD: 5.31 ± 5.01 channels per patient; Fig. 1b). To investigate whether hippocampal ripples are modulated by VSTM, we first extracted ripples from individual hippocampal channels following the well-established protocols in previous studies (Norman et al., 2021, 2019; Sakon and Kahana, 2022). Specifically, raw iEEG data were bipolar re-referenced and filtered between 70 and 180 Hz (see Methods). Then, the amplitude of the data was computed using a Hilbert transform, which was further rectified, squared, smoothed, and converted to z-scores. Ripple events were defined as transient amplitude fluctuations exceeding 4 SD above the pre-encoding baseline (i.e., 200-800 ms before stimulus onset), with durations in the range of 20 − 200 ms (Fig. 2a, see Methods).

**Figure 2.**
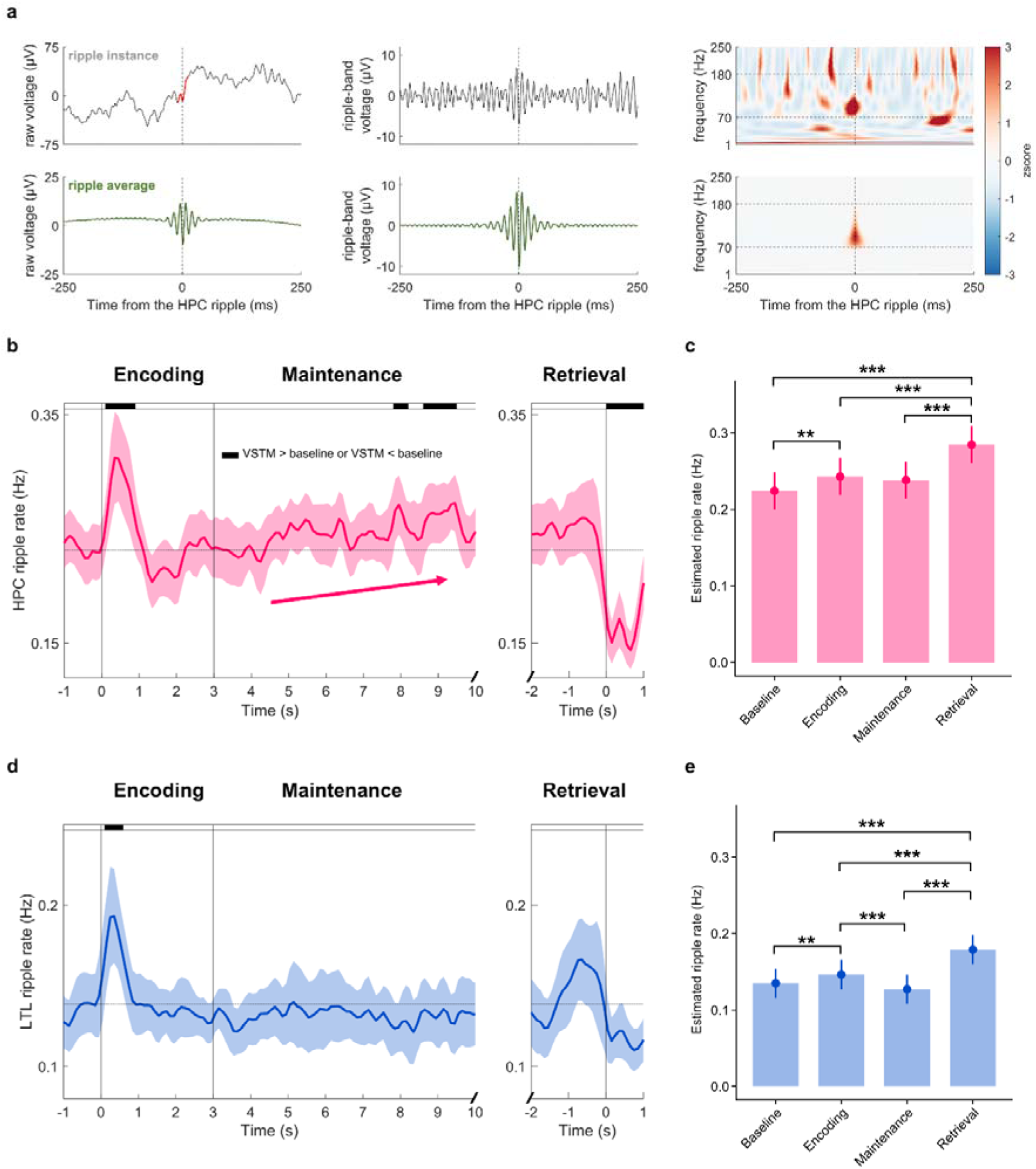
VSTM task-induced ripple rate dynamics. **(a)** Examples of ripple activities from one hippocampal (HPC) channel. Upper (from left to right): a peri-ripple raw iEEG; ripple bandpass (70-180 Hz) filtered iEEG; Power spectrum of the peri-ripple iEEG. Lower (from left to right): averaged peri-ripple raw iEEG across ripples in a channel; averaged ripple bandpass filtered iEEG; averaged peri-ripple power spectrum; **(b)** Time-resolved hippocampal ripple rates averaged across participants and channels during encoding, maintenance, and retrieval. The shaded areas around the lines indicate the standard error of the mean (± 1 SEM). The three vertical lines from left to right indicate onsets of encoding, maintenance, and retrieval, respectively. The peak arrow indicates the temporal ramping-up of ripple rates during the maintenance stage. The bolded grey curve represents the probe onset of each trial during retrieval; Black horizontal bars on the top of the lower panel indicate time clusters where ripple rates differ significantly from the pre-encoding baseline (*p*_cluster_ <□0.05). **(c)** Linear mixed-effect model (LMM) estimated hippocampal ripple rates for pre-encoding baseline (0.2 to 0.8 s before stimulus) and individual task stages. Encoding: 0-3 s; maintenance: 3-10 s; retrieval: probe onset to response. **(d)** Time-resolved LTL ripple rates averaged across participants and channels during encoding, maintenance, and retrieval. **(e)** LMM estimated LTL ripple rates for baseline and individual task stages. Black asterisks indicate significantly higher ripple rates relative to the pre-encoding baseline or significant differences between task stages. *: *p*_FDR_ < 0.05, **: *p*_FDR_ < 0.01, ***: *p*_FDR_ < 0.001, *ns*: not significant.

To validate our detected ripples, we performed peak detection in the unfiltered raw iEEG data and found that 89.84 ± 3.31% of inter-peak intervals (IPIs) within the 70-180 Hz ripple frequency band, confirming that the detected ripples exhibited multi-peak oscillatory waveforms (Buzsáki et al., 1992) (see Fig. S1). We then computed the ripple rate (i.e., number of ripple events per second) for individual hippocampal channels. Across all task stages and participants, the mean ripple rate was 0.29 event/sec (Hz) (see Fig. S1), in line with previous studies (Norman et al., 2021, 2019; Sakon and Kahana, 2022). Moreover, we replicated the novelty effect on ripple rates during memory encoding (Norman et al., 2019). Specifically, hippocampal ripple rates were significantly higher for novel trials (i.e., the first presentation of VSTM items) than repeated trials (i.e., the second and third presentations of VSTM items) during encoding (*p*_cluster_ < 0.001, corrected by the non-parametric cluster-based permutation tests, see Fig. S2).

We next examined ripple rate change across different stages—encoding, maintenance, and retrieval. Hippocampal ripples occurred in 54.26% (SD = 13.55%) of trials during encoding, 78.74% (SD = 13.25%) during maintenance, and 26.85% (SD = 11.27%) during the pre-retrieval stage. Ripple rates were first averaged within each stage per trial. To account for trial-level variability, we then analyzed ripple rate changes across stages using a linear mixed-effects model with task stage as a fixed effect, with participant and trial as random effects. All trials were included in this analysis to ensure consistent trial number and baselines across all stages (see Methods and see Fig. S3 for all hippocampal ripple events for each participant). Our results showed that, compared to the pre-encoding baseline (i.e., −800 to −200 ms relative to stimulus onset), hippocampal ripple rates significantly increased during encoding and retrieval (all *p*s_FDR_ < 0.009, multiple comparisons corrected by false discovery rate (FDR)) but not during maintenance (β = 0.141, *z* = 2.462, *p*_FDR_ = 0.079, Fig. 2c). Direct comparisons showed higher ripple rates during retrieval than during encoding (β = 0.042, *z* = 7.203, *p*_FDR_ < 0.001) and maintenance (β = 0.046, *z* = 8.018, *p*_FDR_ < 0.001), whereas encoding and maintenance did not differ significantly (β = 0.005, *z* = 0.822, *p*_FDR_ = 0.844).

To further examine ripple dynamics at a finer time scale, we performed a time-resolved analysis by calculating the hippocampal ripple rates within consecutive 100 ms non-overlapping sliding time windows for each task stage, averaging across channels for each participant. These ripple rates were then compared to the pre-encoding baseline across participants using cluster-based permutation tests (Maris and Oostenveld, 2007). The analysis revealed significant clusters showing increased hippocampal ripple rates during early encoding (0.15-0.85 s post stimulus-onset, *p*_cluster_ = 0.018; Fig. 2b) and late maintenance (7.85-8.15 s post stimulus-onset and 8.65-9.45 s post-stimulus onset, *p*s_cluster_ < 0.034). In contrast, ripple rates decreased significantly immediately after the response (0.05 s to 1.0 s post-retrieval response, *p*_cluster_ = 0.001), consistent with the decrease observed following long-term memory retrieval in previous studies (Norman et al., 2021; Sakon and Kahana, 2022).

We next performed the same analyses on iEEG data from the lateral temporal lobe (LTL) in the same participants (132 channels; mean ± SD: 10.15 ± 7.82 channels per patient; Fig. 1b). On average, LTL ripples occurred in 32.17% (SD = 14.69%) of trials during encoding, 52.34% (SD = 17.82%) during maintenance, and 14.99% (SD = 10.30%) during the pre-retrieval stage. Similar to the hippocampus, LTL ripple rates during encoding and retrieval were significantly higher than the pre-encoding baseline (both *ps*_FDR_ < 0.002), whereas maintenance did not differ from baseline (β = −0.008, *z* = −2.505, *p*_FDR_ = 0.059, Fig. 2d-e). Ripple rates during both encoding and retrieval were greater than during maintenance (both *p*s_FDR_ < 0.001), and retrieval showed higher ripple rates than encoding (β = 0.033, *z* = 10.436, *p*_FDR_ < 0.001). The time-resolved group-level analysis revealed a significant cluster of increased ripple activity during early encoding (0.15-0.55 s post-stimulus onset, *p*_cluster_ = 0.047; Fig. 2d). Given that both the hippocampal and LTL ripple rates increased within the 0-1 s of encoding, we re-performed the comparisons of the averaged ripple rates between task stages using this 0-1 s encoding time window while keeping other stages unchanged, and the pattern of the main results remains (see Fig. S4).

To assess whether hippocampal ripple rates were associated with VSTM performance, we first compared ripple rates between remembered and forgotten trials (i.e., memory accuracy), as well as between fast and slow responses among remembered trials (i.e., retrieval speed). Replicating previous studies (Sakon and Kahana, 2022), hippocampal ripple rates are higher for successful than unsuccessful long-term memory retrieval (Fig. S2). However, no significant differences were observed between remembered and forgotten trials for any VSTM stages (all *p*s_cluster_ > 0.466; Fig. S2). Besides, fast and slow trials were defined as lower or higher than each participant’s median reaction time (RT), respectively. Faster trials exhibited significantly higher ripple rates than slow trials during the pre-response retrieval (*p*_cluster_ = 0.014; Fig. S2). The same analyses on LTL channels revealed no significant effects for either memory accuracy or retrieval speed (*ps*_cluster_ > 0.237, Fig. S2).

These findings together suggest that while both hippocampal and LTL ripple rates were modulated by the VSTM task, these ripple rates were not associated with VSTM performance. Besides, for the hippocampal channels, ripple rates were above pre-encoding baseline during the late maintenance time windows, suggesting a potential ramping-up pattern that was tested in the following analyses.

### Ramping-up hippocampal ripples during maintenance supports successful VSTM

Next, we tested whether hippocampal ripples ramp up during the maintenance stage, as proposed by the dynamic coding framework (Stokes, 2015). Using a generalized linear mixed-effects model (GLMM) with ripple occurrence as the dependent variable, maintenance time as fixed effects, and random intercepts/slopes for participants and trials, we observed a significant fixed effect of time across all trials (β = 0.012, *z* = 4.678, *p* < 0.001, Fig. S5), suggesting a ramping-up effect of hippocampal ripples. A similar analysis of the broadband high-frequency activity (HFA) (70-180 Hz) during maintenance revealed a non-significant ramping-up trend (*p* = 0.074), and the ramping slopes did not significantly differ between ripple events and HFA (*p* = 0.662).

To examine whether this ramping-up predicted the VSTM performance, we further fitted a similar GLMM model with the interaction term: maintenance time X VSTM accuracy (i.e., remember versus forget) as fixed effects, and all trials were included in this analysis (see Methods). Our results revealed a significant interaction effect between VSTM accuracy and maintenance time (β = 0.018, *z* = 2.171, *p* = 0.030, Fig. 3a), indicating that the ripple ramping-up effects differed between remembered and forgotten trials. Post hoc analyses revealed a significant ramping-up for remembered trials (β = 0.014, *z* = 5.024, *p*_FDR_ < 0.001), but not for the forgotten items (β = −0.002, *z* = −0.282, *p*_FDR_ = 0.778).

**Figure 3.**
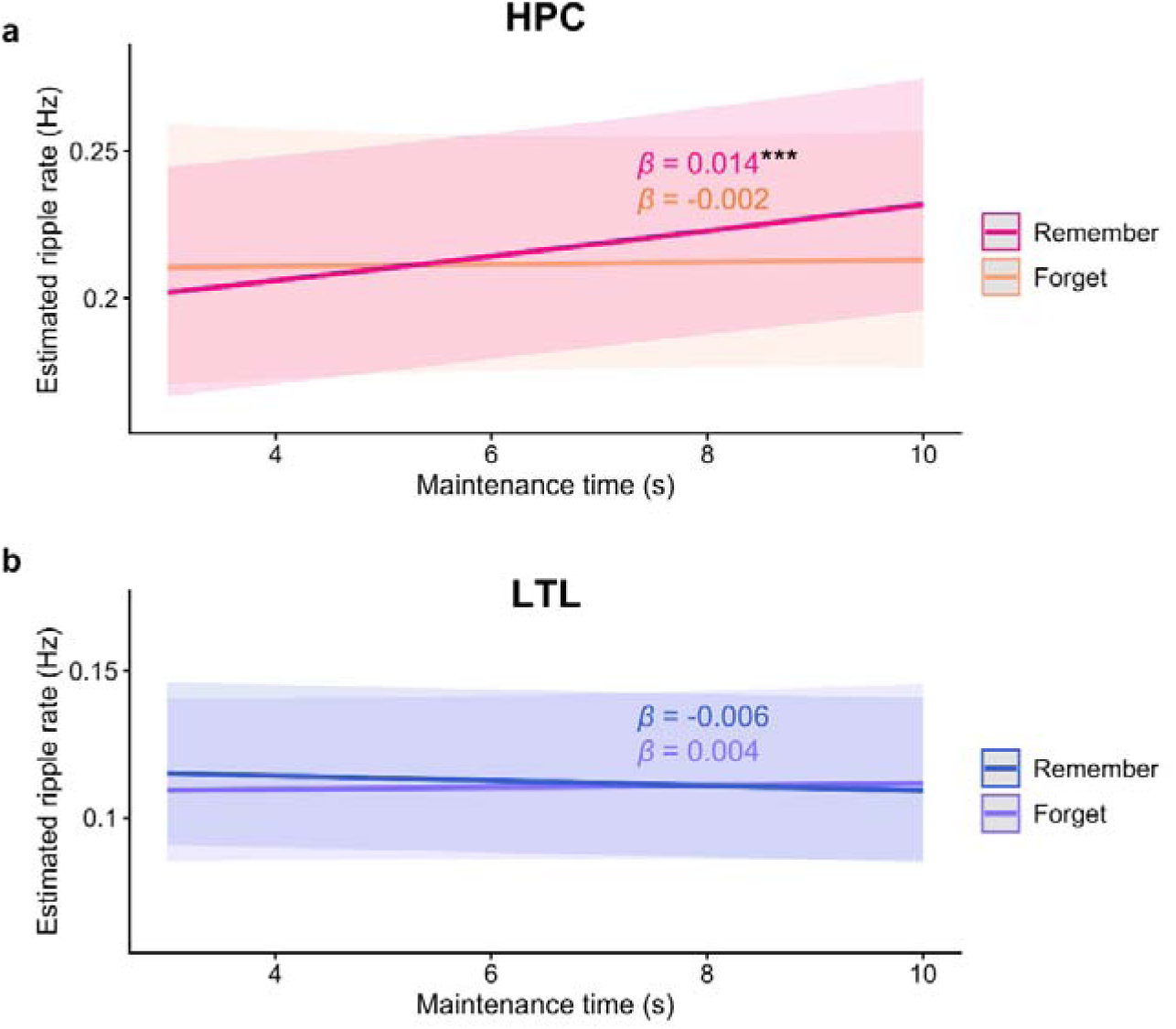
Ripple ramping-up effects during maintenance. **(a)** Hippocampal (HPC) ripple ramping-up effects for remembered vs. forgotten trials. **(b)** Lateral temporal lobe (LTL) ripple ramping-up effects for remembered vs. forgotten trials. The shaded areas around the lines indicate ± 1 SEM. β: estimated fixed effect coefficients for remember or forget conditions. ***: *p*_FDR_ < 0.001.

Further control analyses revealed that the VSTM accuracy-related hippocampal ramping-up effect was robust across alternative ripple definitions (e.g., ripple duration ≥ 25 ms or alternative frequency bands: 80-120 Hz and thresholds following previous research (Vaz et al., 2019; see Methods and Fig. S5) and remained significant using conventional group-level analysis (see Methods and Fig. S5; see also Table S1 for participant-level ramping-up slopes). Moreover, the hippocampal ramping-up effect was unrelated to retrieval speed (β = 0.009, *z* = 1.570, *p* = 0.117; Fig. S5) and did not differ between novel and repeated trials (β = −0.009, *z* = −1.716, *p* = 0.086; Fig. S5).

We also tested the ripple ramping-up during maintenance in the LTL. Unlike the hippocampus, no significant ramping-up was observed (β = −0.005, *z* = −1.772, *p* = 0.076, see Fig. S5), nor was the change in ripple rates over maintenance associated with VSTM accuracy (time × accuracy interaction effect: β = −0.010, *z* = −1.246, *p* = 0.213; remember: β = −0.006, *z* = −2.033, *p_FDR_* = 0.084; forget: β = 0.004, *z* = 0.507, *p_FDR_* = 0.612; Fig. 3b) or RT (time × RT interaction effect: β = 0.006, *z* = 1.087, *p* = 0.277, see also Fig. S5). To ensure that the absence of a ramping-up effect was not due to low signal-to-noise channels, we restricted the analysis to 89 bipolar channel pairs with at least one contact in LTL gray matter or both contacts within 2 mm of it, and the results remained (see Fig. S5). In addition, the absence of the ramping-up effect in the LTL cannot be attributed to greater signal heterogeneity compared with the hippocampus (see Fig. S5). Moreover, a direct comparison between the ramping-up effects of the HPC versus the LTL revealed a significant three way time × accuracy × region interaction effect (β = 0.053, *z* = 2.012, *p* = 0.044), supporting the dissociation of the ripple ramping-up patterns between the HPC and the LTL (see Methods).

### HPC-LTL coupled ripples associate with VSTM performance

Previous research suggested that hippocampal ripples support memory via hippocampal-neocortical interactions (Rothschild et al., 2017), which may be supported by ripple couplings across regions (Khodagholy et al., 2017; Vaz et al., 2019; Verzhbinsky et al., 2024). We therefore examined whether HPC-LTL ripple coupling supports VSTM. Following previous research (Dickey et al., 2022), we first computed the LTL ripple rate within a broader window of ± 0.5 s of each hippocampal ripple peak (Fig. 4a; see Methods) across all task stages as well as within each task stage. This broader window was employed to examine the lead-lag between ripples of these two regions, and this analysis was performed exclusively among trials in which the HPC ripples were detected. We then computed LTL ripple rates within ± 0.5 s of randomly selected, hippocampal ripple-free time points 1000 times, resulting in a surrogate distribution. The empirical coupled ripple rates were then z-scored against the surrogate distribution to yield normalized LTL ripple rates, which were coupled to the HPC ripple peaks (see Methods). Our results revealed that the normalized LTL ripple rates were significantly greater than zero across all task stages (*p*_cluster_ < 0.011; Fig. 4b), and within individual stages (all *ps*_cluster_ < 0.047). In addition, peak-to-peak lag analysis revealed no significant difference between LTL and HPC ripple peaks in any stage (*p*s_FDR_ > 0.126).

**Figure 4.**
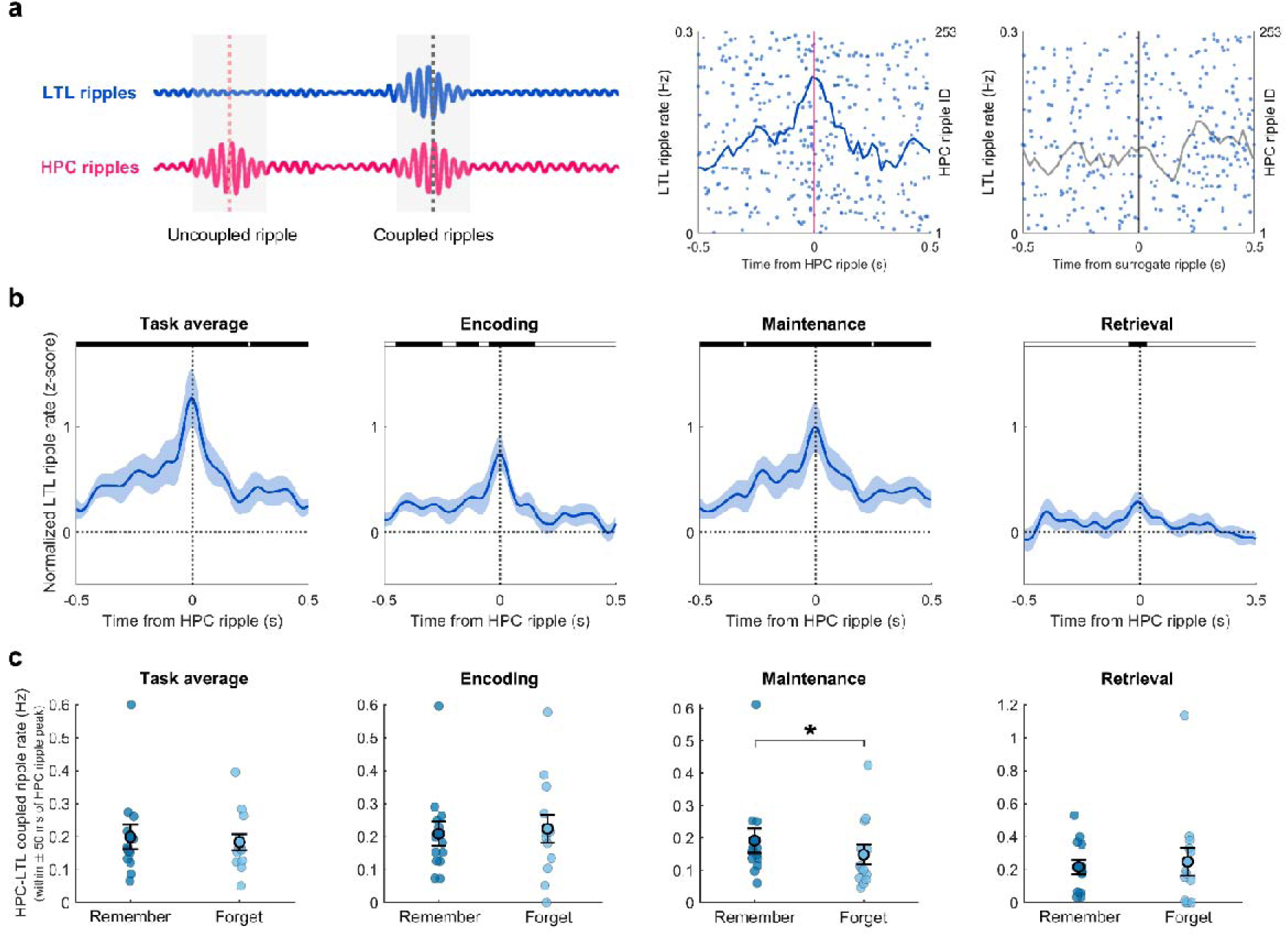
HPC-LTL coupled ripples. **(a)** Left: Illustration of coupled ripples between HPC and LTL (second shaded area) and uncoupled ripples (first shaded area). Middle: LTL ripple rates time-locked to an exemplar HPC ripple from one participant; Right: LTL ripple rates time-locked to surrogate time points without HPC ripples from the same channels. Each row indicates LTL ripples locked to a single HPC ripple peak or surrogate time point. Each blue dot represents an LTL ripple, and the curve shows LTL ripple rates across all trials surrounding HPC ripple peaks or surrogate time points. **(b)** Normalized LTL ripple rates locked to the HPC ripple peak (i.e., time 0 on the x-axis) across all task stages (i.e., task average) and within individual task stages. Black bars at the top indicate time windows with significant differences between conditions (survived after cluster-based permutation tests: *p*_cluster_ <□0.05). The shaded areas around the lines indicate ± 1 SEM. **(c)** HPC-LTL coupled ripple rate (i.e., LTL ripple rates occurred within ± 50 ms of the HPC ripple peak) for VSTM remembered versus forgotten trials. *: *p*_FDR_ < 0.05.

We next examined whether HPC-LTL coupled ripples contribute to VSTM performance. Notably, the coupled ripples were defined as LTL ripples occurring within ± 50 ms of HPC ripple peaks following previous research (Vaz et al., 2019). We then compared the coupled ripple rates between remembered and forgotten trials. The results revealed significantly higher coupled ripple rates in remembered trials compared to forgotten trials during maintenance (*t*(12) = 2.897, *p*_FDR_ = 0.040; Fig. 4c). This effect remained significant after matching trial counts between remember and forget conditions using bootstrapping (*p* = 0.017, see Methods). In contrast, no significant differences were observed between remembered versus forgotten trials during encoding or retrieval stages (*p*s_FDR_ > 0.726), and the interaction effect between stages and memory accuracy was not significant (*F*(2,24) = 0.49, *p* = 0.619). In addition, peak-to-peak lag analysis among remembered trials revealed no significant difference between LTL and HPC ripple peaks for the coupled ripples in any task stage (*p*s_FDR_ > 0.069). Together, these findings suggest that HPC-LTL ripple coupling during the maintenance was associated with successful VSTM.

We next examined the temporal dynamics of coupled ripples during maintenance. HPC-LTL coupled ripples showed non-significant ramping patterns for both remembered and forgotten trials (*p*s_FDR_ > 0.281), with no significant difference between these two conditions (*p* = 0.506; see Fig. S6).

### HPC-LTL coupled ripples coordinate memory reactivation in the LTL

We next tested whether HPC-LTL coupled ripples support the VSTM reactivation in the LTL. To this end, we applied a multivariate decoding method to the spectral power of LTL channels for individual participants to classify categories of learned pictures (see Methods). Classifiers were trained on each encoding time window and tested across encoding, maintenance, and retrieval time windows. Among remembered trials, we identified significant clusters showing decoding accuracies above chance level (25%) during encoding, early maintenance, and retrieval (all *ps*_cluster_ < 0.039, Fig. 5a). When averaged across all time windows within each stage, decoding accuracy remained significantly above chance for individual stages (all *ps*_FDR_ < 0.003, Fig. 5b). Note that the above-chance decoding during the maintenance stage cannot be simply attributed to the lingering sensory input immediately following encoding. This effect remains robust in the 3-4 s post-encoding interval (Fig. S7). Furthermore, remembered items showed greater decoding accuracy than forgotten ones during the pre-response retrieval stage (Fig. S7). These results suggested that VSTM is represented and reactivated in the LTL.

**Figure 5.**
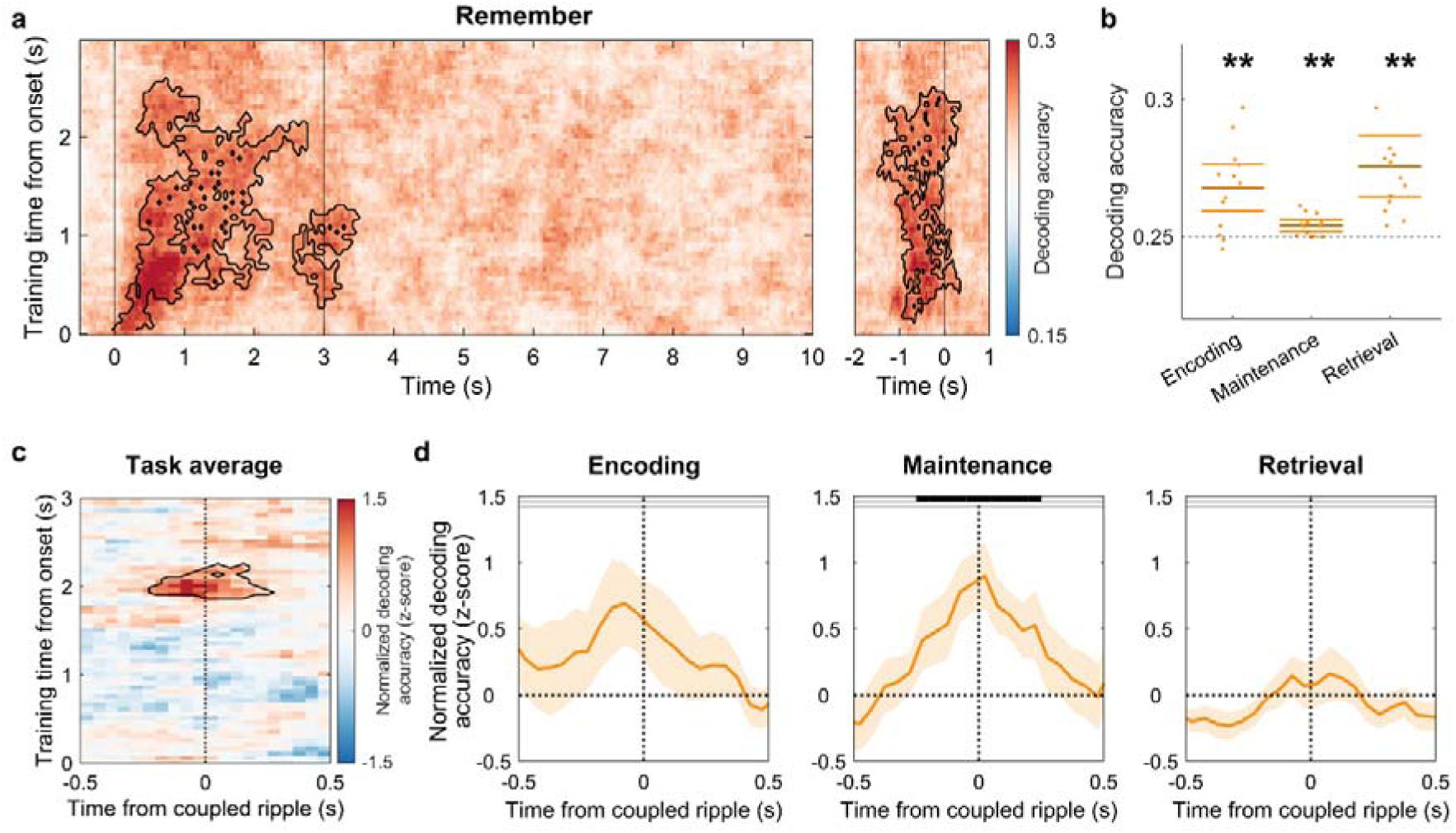
Coupled ripples coordinate memory reactivation in the LTL. **(a)** LTL decoding accuracy of remembered trials compared to chance level (0.25) across the task (left: encoding and maintenance, 0 indicates stimulus onset; right: retrieval, 0 indicates behavior response). Clusters with significantly above-chance decoding accuracy (survived cluster-based permutation test) are circled by black lines. **(b)** Decoding accuracies during encoding, maintenance, and retrieval stages are significantly above chance. **(c)** Normalized decoding accuracy time-locked to HPC-LTL coupled ripples relative to surrogate distribution. The black-circled cluster indicates normalized decoding accuracy significantly above zero. **(d)** Coupled ripple-locked normalized decoding accuracy averaged across the late encoding cluster identified in (c). Black bars at the top indicate significant clusters with normalized decoding accuracy significantly above zero. All clusters survived cluster-based permutation tests (*p*_cluster_ < 0.05). The shaded areas around the lines indicate the ±1 SEM. **: *p*_FDR_ < 0.01.

We then tested whether this reactivation was temporally locked to the HPC-LTL coupled ripples during remembered trials. To enhance signal-to-noise ratio, we pooled HPC-LTL coupled ripples across all task stages (mean ± SD: 439 ± 125 coupled ripples per participant) and aligned decoding accuracy to ± 0.5 s around hippocampal ripple peaks of the LTL coupled ripples. Notably, the decoding accuracy was z-scored relative to the surrogate distribution, where decoding accuracy aligned to non-coupled ripple time points, and then tested against zero (see Methods). Our results revealed a significant positive cluster when the classifier was trained on late encoding windows (1.85-2.25 s post-stimulus; *p*_cluster_ = 0.029; Fig. 5c), indicating LTL memory reactivation time-locked to coupled ripples. Further analysis based on the identified cluster, we found that this coupled ripple-locked decoding accuracy was significant during maintenance (*p*_cluster_ = 0.004, Fig. 5d), but not during the encoding or retrieval stage (*p*s_cluster_ > 0.118). Further analysis revealed that coupled ripple-locked decoding accuracy was higher during maintenance than during the retrieval stage (*p*_cluster_ = 0.024), but not significantly different from that during the encoding stage (*p*s_cluster_ > 0.800) (see Fig. S8). Notably, coupled ripple-locked LTL decoding accuracy in the clusters in Fig. 5c and 5d was significantly above chance (i.e., 25%; all *p*s < 0.002).

As a control analysis, we also examined memory reactivation in the hippocampus and its temporal link with HPC-LTL coupled ripples. Although hippocampal decoding accuracy was above chance when averaged across time windows for each task stage (*ps*_FDR_ < 0.031), it was not significantly time-locked to HPC-LTL coupled ripples (*ps*_cluster_ > 0.470; Fig. S8). Further control analyses to test the specificity of coupled ripple revealed that decoding accuracy in neither the hippocampus nor the LTL was modulated by independent, uncoupled ripples (i.e., hippocampus or LTL ripples without coupling) (*ps*_cluster_ > 0.203; Fig. S8). These findings suggest that coupled HPC-LTL ripples coordinate memory reactivation in the LTL, rather than in the hippocampus, during VSTM maintenance.

We also examined the temporal dynamics of coupled ripple-locked memory reactivation during maintenance. The memory decoding accuracy showed a significant time × VSTM accuracy interaction (*p* = 0.009; see Fig. S6), with a significant decline on forgotten trials (*p*_FDR_ = 0.031), while relatively stable on remembered trials (*p*_FDR_ = 0.214). Within remembered trials, coupled ripple-locked memory decoding accuracy showed a non-significant numeric increase across the maintenance stage (*p* = 0.586; see Fig. S6).

### VSTM-related ripple dynamics are not associated with subsequent long-term memory performance

To rule out potential confounds from long-term memory (LTM) formation, we separated VSTM-remembered trials based on whether they were later remembered or forgotten in the LTM test, as well as compared the hippocampal ramping-up effect, HPC-LTL coupled ripples, and coupled ripple-locked memory reactivation between these two conditions. Our results showed that the hippocampal ripple ramping-up effect was significant for both subsequently LTM remembered (β = 0.017, *z* = 4.141, *p_FDR_* < 0.001) and forgotten trials (β = 0.011, *z* = 2.987, *p_FDR_* = 0.003; see Fig. S9). Critically, no significant difference was observed between subsequently LTM remembered and forgotten items (β = 0.003, *z* = 0.439, *p* = 0.661). A further analysis examining the three-way interaction between time, memory accuracy, and test type (VSTM vs. LTM) for hippocampal ripples revealed a trend but statistically non-significant interaction effect (β = 0.039, *z* = 1.809, *p* = 0.071). Moreover, neither the coupled ripple rate nor ripple-locked reactivation differed between subsequently LTM remembered and forgotten trials (Fig. S9). Together, these analyses suggest that our main findings reflect mechanisms underlying VSTM rather than LTM.

## Discussion

Previous rodent and recent human studies have implicated hippocampal ripples in long-term memory consolidation and retrieval (Joo and Frank, 2018; Norman et al., 2019). Our study extends this field by showing that hippocampal ripples also contribute to VSTM maintenance. We found that hippocampal ripples progressively ramped up during the maintenance period, and this ramping-up effect was associated with successful VSTM. Moreover, hippocampal ripples were temporally coupled with LTL ripples, and these coupled ripples coincided with memory reactivation in the LTL during maintenance.

The hippocampal ripple rate progressively ramps up over the 7-s maintenance period, while the overall average rate during this period does not exceed the pre-encoding baseline. This dynamic profile aligns directly with the dynamic coding framework of VSTM (Stokes, 2015). The ramping-up in hippocampal ripple rates is associated with memory accuracy but not reaction time, suggesting its role in proactive memory retrieval during VSTM rather than general arousal or motor preparation (Kikumoto et al., 2022; van Ede et al., 2019). Moreover, this ramping-up effect is unlikely to reflect signal drifts over long intervals, since it was observed exclusively in remembered trials and has also been reported in studies using shorter retention periods, such as 1.5 s (Lundqvist et al., 2016) or 3 s (Murray et al., 2017). Nonetheless, we cannot rule out the possibility that the magnitude of this effect may be affected by the duration of the delay period. Future studies could systematically manipulate retention intervals to test this possibility.

However, neither the coupled ripples nor the coupled ripple-locked memory reactivation shows a significant ramping-up effect. Nevertheless, our results indeed observed that, in contrast to the significant decrease in decoding accuracy for forgotten trials, the remembered trials showed relatively stable memory reactivation during the maintenance period, suggesting that coupled ripples may coordinate discrete reactivation events in supporting the stable VSTM maintenance. Meanwhile, we observed that hippocampal ripple rates decreased immediately following the response, a pattern consistent with that during episodic memory retrieval (Norman et al., 2021; Sakon and Kahana, 2022), which may reflect diminished hippocampal engagement following memory retrieval.

While the ramping-up hippocampal ripples align with the dynamic coding framework of VSTM, they appear to contrast with prior reports of persistent hippocampal spiking during short-term memory maintenance (Kamiński et al., 2017; Kornblith et al., 2017). Several factors may account for this discrepancy, including differences in neural measures and task demands. First, prior studies identifying persistent activity were mostly based on single-unit recordings (Fuster and Alexander, 1971; Kamiński et al., 2017; Kornblith et al., 2017), revealing sustained firing in a minority of stimulus-selective neurons. In contrast, our local field potential (LFP) recordings may reflect a dynamic coding scheme at the population level (Stokes, 2015). Compared to persistent firing, this dynamic population coding is less constrained by memory capacity limits and may offer more flexible control while minimizing inter-item competition (Lundqvist et al., 2018a). Second, contemporary short-term memory models suggest that items within the focus of attention are maintained via persistent activity, while unattended items are stored in an activity-silent state (Cowan, 2019; Trübutschek et al., 2017). Therefore, multi-item VSTM tasks with higher attentional demands may favor persistent activity, while our single-item task may have allowed memory to drift out of focus, favoring an overall activity-silent state with discrete reactivation bursts. Corroborating this possibility, a previous study found persistently increased hippocampal gamma band activity during multi-item working memory maintenance but not for the single-item (Axmacher et al., 2007).

In addition, our results indicate that hippocampal ripples are unlikely to be the sole mechanism supporting VSTM. While we validated the ripple events, our current data did not show significant differences in the ramping-up effects between hippocampal ripples and HFA. Recent research demonstrated that HFA exhibited broad subsequent memory effects across MTL regions, whereas hippocampal ripples were more selectively associated with subsequent temporal and semantic clustering during recall (Sakon and Kahana, 2022). Thus, future studies with denser MTL coverage and direct comparisons between ripples and HFA under specific task manipulations are needed to better investigate whether ripples make a functional contribution beyond general high-frequency activation. Moreover, prior work has shown that gamma bursts in the prefrontal cortex gate access to encoded VSTM representations and that ramping of such bursts supports working memory in primates (Lundqvist et al., 2016). Recent rodent research demonstrates that replay sequences can occur even in the absence of sharp-wave ripples (Widloski and Foster, 2025). These findings suggest that hippocampal ripple ramping-up should be interpreted as one component within a broader, distributed set of dynamic mechanisms supporting short-term memory maintenance.

Our findings further demonstrated that ripple coupling strength during maintenance was associated with VSTM accuracy, which is consistent with prior findings showing that increased hippocampal-neocortical functional connectivity is associated with subsequent memory performance(de Voogd et al., 2016). Comparing with previous research emphasizing low-frequency coordination supporting VSTM (Liu et al., 2020), our findings suggest hippocampal-cortical communication could be achieved through an alternative mechanism: the co-occurrence of ripples. However, our ripple peak lag analysis revealed little evidence regarding the directionality of coupled ripples during the maintenance period. Recent evidence shows that ripples in different brain areas often co-occur with near-zero phase lag, providing a temporally precise mechanism for efficient inter-regional communication (Verzhbinsky et al., 2024). Such synchronous coupled ripples may reflect a shared or rapidly reciprocal activation between hippocampus and neocortex, rather than a unidirectional drive. Although the absence of significant directionality does not necessarily undermine the functional relevance of coupled ripples, it leaves the open question of whether hippocampal ripples actively gate cortical reactivation or merely co-occur with it. Future studies with directed connectivity manipulations or measures will be necessary to resolve this issue.

Despite the observed HPC-LTL coupled ripples and the role of LTL in representing VSTM content, we found the LTL ripple rates did not ramp over the maintenance. This absence of ripple ramping-up in the LTL channels cannot be attributed to greater ripple heterogeneity since controlling the ripple-rate variability did not affect the results (Fig. S5). There are at least two possibilities: First, hippocampal ripple coupling selectively influence a subset of LTL ripple events, given the coupled ripples showing a ramping-up trend, while uncoupled ripple decreasing overing the maintenance period (see Fig. S10); Second, LTL ripple dynamics are shaped by broader cortical inputs beyond hippocampal coupling alone, as prior work indicates that LTL activity can be strongly influenced by prefrontal and parietal inputs (Axmacher et al., 2008; Hirabayashi et al., 2024; Xu, 2023). Moreover, the absence of LTL ramping-up does not necessarily contradict the dynamic coding framework, as the region maintaining decodable content and the region exhibiting ramping dynamics may differ yet interact to support VSTM maintenance. Due to electrode coverage limitations, we cannot evaluate other key cortical areas, such as the frontal or parietal cortex, where VSTM information is traditionally maintained (Bettencourt and Xu, 2016; Kamiński et al., 2017). To further test these possibilities, future studies with broader cortical sampling and higher ripple counts will be necessary.

Our results showed that decoding accuracy during maintenance was locked to HPC-LTL coupled ripples, rather than isolated ripples in either the hippocampus or the LTL, highlighting the necessity of hippocampal-neocortical communication in supporting memory representational reactivation. Notably, while encoding ripples peaked early after stimulus onset, the classifier trained on late encoding activity showed the strongest ripple-locked reactivation during maintenance. This temporal discrepancy is consistent with the memory representational transformation framework, which proposes that mnemonic representations evolve over time (Xue, 2022). Early encoding activity may primarily reflect low-level visual/perceptual features, whereas later encoding may contain more stable category-level or semantic representations. Prior intracranial and neuroimaging studies have implicated lateral temporal regions in such higher-level coding (Ralph et al., 2017). Given that our decoder targeted category-level information, it is reasonable that the decoder performed well when trained on this later window. Thus, delay-period hippocampal-LTL coupled ripples may preferentially reinstate higher-level category/semantic representations during VSTM maintenance. Early perceptual representations may instead rely more on hippocampal-visual cortical interactions, which could not be assessed due to limited occipital coverage. In addition, no significant coupled ripple-locked decoding accuracy was observed during encoding or retrieval. This likely arises because encoding and retrieval involve salient visual inputs (target picture or lure), such that the LTL decoding accuracy is probably driven by feedforward perceptual processing. In contrast, during maintenance, where no external input is available, coupled ripples may serve as a specialized, internally generated mechanism to transiently reactivate and sustain representations.

Prior work suggested that short-term performance can be supported by pre-existing semantic knowledge and episodic experience, such that hippocampal engagement may reflect interactions between visual perceptual input and long-term pre-existing knowledge/episodes, rather than a strictly process-pure visual short-term memory (VSTM) (Chung et al., 2024; Cowan, 2001; Jeneson and Squire, 2012). However, recent evidence suggests that simplified stimuli may underestimate how the brain maintains meaningful information in real-world contexts (Chung et al., 2024). Indeed, semantically rich stimuli can enhance working memory capacity by enabling access to long-term representations (Brady et al., 2016; Brady and Störmer, 2022; Thibeault et al., 2024; Yu et al., 2025). We therefore interpret our findings as reflecting mechanisms that support the short-term maintenance of meaningful, real-world visual representations. Future studies aiming to more strictly dissociate VSTM from LTM contributions should employ artificial or abstract stimuli that minimize pre-existing knowledge associations.

In addition, our task involves cue-picture associations, which makes it challenging to fully disentangle VSTM from LTM-related processes. Nevertheless, our findings showed that both the ramping-up hippocampal ripple rates and the HPC-LTL coupled ripples during maintenance were prominently associated with VSTM accuracy but not with subsequent LTM accuracy. These findings suggest that the ripple dynamics we observed primarily support the short-term maintenance of complex visual features and their binding within the pictures themselves rather than the formation of long-term memory for cue-picture pairs, which has been shown in previous research (Pacheco Estefan et al., 2019). This interpretation is strengthened by recent studies showing that hippocampal ripples and hippocampal neuronal firing support VSTM of complex naturalistic pictures in the absence of subsequent LTM tests or cue-picture associations (Daume et al., 2024; Verzhbinsky et al., 2025). Moreover, converging evidence from studies on neural oscillations, patients with focal hippocampal damage, and neural modulation demonstrates that the hippocampus plays a critical role in VSTM, where high-resolution feature binding is required (Borders et al., 2022; Koen et al., 2017; Yonelinas, 2013).

Beyond binding demands, the hippocampus also contributes to VSTM for simple visual features that minimally engage LTM, such as color squares, when memory precision is dissociated from binding errors or when relational demands are minimized (Borders et al., 2022; Choo et al., 2025). For instance, previous research found that patients with hippocampal damage exhibited reduced precision for simple color memories after brief delays, yet showed no increase in relational binding errors (Borders et al., 2022). Similarly, a recent study demonstrated that MTL lesions selectively impair the precision of VSTM representations rather than the quantity of items retained (Choo et al., 2025). In addition, the hippocampus is also known to engage in tasks with relatively low memory precision demands, such as change detection paradigms requiring only coarse-level discrimination between targets and distinct lures (Daume et al., 2024; Kamiński et al., 2017; Ranganath and D’Esposito, 2001). These findings collectively indicate that the hippocampus supports a wide range of VSTM tasks. Future work should systematically examine whether key factors, such as relational binding demands and memory precision, shape hippocampal ripple dynamics, using experimental manipulations or multi-component modeling (Hitch et al., 2025).

Notably, the absence of the hippocampal ramping-up effect during maintenance in predicting LTM does not conflict with the established role of hippocampal ripples in long-term memory. Converging evidence suggests that ripple contributions to LTM are most evident during offline consolidation windows, including sleep and post-encoding rest, rather than during online task execution. In rodents, suppressing post-learning sleep ripples impairs subsequent memory (Girardeau et al., 2009). In humans, ripple-coupled replay during NREM sleep predicts later memory success (Zhang et al., 2018), and post-encoding awake-rest ripple activity, rather than encoding-period activity, is associated with subsequent memory performance (Zhang et al., 2024). Thus, the lack of association between VSTM-delay ripples and later cued recall raises a possibility: ramping-up ripple dynamics during active maintenance may support online VSTM, whereas LTM formation may depend more on ripple events occurring during post-encoding time windows. Our study cannot rule out the possibility that any relationship between VSTM delay-period ripples and later LTM was obscured by intervening processes between maintenance and recall, including interference resistance, offline consolidation during rest, and retrieval-related factors. Future studies incorporating parametric manipulations, such as varying memory load, delay duration, or lure similarity, will be essential to directly test whether hippocampal ripple activity selectively scales with VSTM demands, thereby providing stronger evidence for its specific role in online maintenance versus LTM-related processes.

To conclude, our study provides direct electrophysiological evidence that hippocampal ripples—and their coordination with neocortical regions—support VSTM. We show that ripple activity in the hippocampus ramps up during maintenance and is associated with memory accuracy. These findings support a dynamic coding model of VSTM and suggest that hippocampal ripples orchestrate neocortical reactivations that sustain stable VSTM representations. By linking ripple dynamics to both representational reactivation and interregional coupling, our results extend the hippocampus’s role from long-term formation to short-term memory maintenance and offer new insights into the neural mechanisms unifying these memory systems.

## Methods

### Participants

We reanalyzed the data from a previous study (Liu et al., 2020). We included thirteen epilepsy patients (mean age ± SD: 26.77 ± 5.48 years, 7 females) who had electrodes implanted in both the hippocampus and LTL. The iEEG data were recorded at the Center of Epileptology, Xuanwu Hospital, Capital Medical University, Beijing, China. The study adhered to the latest version of the Declaration of Helsinki and was approved by the Institutional Review Board at Xuanwu Hospital. All participants have a normal or corrected-to-normal visual acuity. They all signed written informed consent before the experiment.

### Stimuli and procedures

The study consisted of a visual short-term memory (VSTM) task, followed by a 1-minute countback task and a short break (1 to 4 minutes), and then a cued-recall long-term memory test. The study used 56 pictures of familiar everyday objects and 112 two-character Chinese verbs. The pictures were drawn from four categories—fruits, animals, electrical devices, and furniture—with 14 images per category. Each picture was randomly paired with two different cue words across two consecutive experimental runs.

The VSTM was assessed using a modified Delayed Match-to-Sample (DMS) task. In each run, participants studied 14 unique word-picture pairs, each repeated three times, resulting in 42 trials per run. Each trial started with a brief fixation period (300 ms), followed by a jittered blank interval (800-1200 ms). A cue word and its associated target picture were then presented centrally for 3 s, during which participants were instructed to encode the association between the word and the picture. Immediately following the encoding stage, the picture disappeared while the cue word remained on screen for 7 s. During this maintenance stage, participants were instructed to mentally maintain a vivid image of the target picture. Displaying the cue word during maintenance served to reinforce the word-picture association, reduce working memory load, and minimize distraction. It was then followed by an immediate retrieval stage, during which a probe picture was displayed on the screen, and participants were instructed to determine whether it matched the target picture by pressing one of two buttons within 2 s. The probe picture was either the same as the target picture (50%, match trials) or a highly similar lure picture (50%, nonmatch trials). Each participant completed between 4 and 8 DMS runs (mean ± SD: 6.14 ± 1.46), resulting in a total of 168-336 trials (mean ± SD: 258.00 ± 61.32) per participant.

Following the completion of each VSTM run, participants performed a 1-minute mathematical countback task, followed by a brief rest period (1-4 min) before the LTM cued-recall test. During each LTM test run, 14 learned, unique cue-picture pairs were tested in a randomized order, with each pair tested only once. There are, on average, 88.31 ± 19.28 trials per participant. For each trial, participants first performed a self-paced, binary keypress to indicate subjective retrieval success (“remember” vs. “don’t remember”). Upon a subjective “remember” response, an objective category verification prompt was presented for a maximum of 3 s, requiring a 4-alternative forced-choice response to identify the semantic category of the associated target picture (animals, electronics, fruits, or furniture). Trials were operationally classified as LTM “remembered” trials only if the subjective recall was accompanied by a correct objective category verification. Trials receiving a “don’t remember” response or an incorrect category report were classified as LTM “forgotten” trials.

### Data recordings and preprocessing

Intracranial EEG data were recorded from depth electrodes, each containing 12 or 16 channels (2 mm in length, 0.8 mm in diameter, spaced 1.5 mm apart). Data were collected using amplifiers from Brain Products (Brain Products GmbH), NeuroScan (Compumedics Limited), or Nicolet (Alliance Biomedica Pvt Ltd.) electroencephalogram systems, with sampling rates of 2,500, 2,000, and 2,048 Hz, respectively. During the online recordings, data from all channels were referenced to a common subcutaneous channel. During the offline preprocessing, we first removed the channels within the epileptic loci and channels that were severely contaminated by epileptic activity. The remaining channels in the hippocampus (HPC) and lateral temporal lobe (LTL) were visually inspected and bipolar referenced to channels on the same electrode. Because not all participants had multiple hippocampal channels on the same electrode, HPC channels were bipolar re-referenced to the nearest white matter channel (Chen et al., 2021; Norman et al., 2021). LTL channels were bipolar re-referenced to an adjacent channel within the LTL (Pacheco Estefan et al., 2019).

### Channel localization

The identification of channel locations included the following steps. First, each participant’s post-implantation CT scan was co-registered with their pre-implantation MRI. The MRI was then normalized to Montreal Neurological Institute (MNI) space using Statistical Parametric Mapping 12 (SPM12). Anatomical localization and 3D visualization of electrode channels were conducted using the 3D Slicer platform (https://www.slicer.org/). To assign anatomical labels, structural MRIs were segmented with FreeSurfer (https://surfer.nmr.mgh.harvard.edu), and the nearest cortical or subcortical label was assigned to each channel. Channels in the hippocampus were further verified through visual inspection in each participant’s native anatomical space.

### Ripple detection and peri-ripple spectral analysis

Ripple detection followed the established procedures outlined in previous studies (Norman et al., 2021, 2019; Sakon and Kahana, 2022). First, a 70-180 Hz bandpass linear-phase Hamming-windowed finite impulse response (FIR) filter was applied to the bipolar re-referenced iEEG data on HPC and LTL channels, with a transition width of 5 Hz. We then computed the analytic signal amplitude (squared) using the Hilbert transform. To determine the ripple detection threshold, we: (1) identified and clipped extreme values (≥4 SD above mean) using the Least-Median-Square (LMS) method to reduce outlier bias; (2) smoothed the clipped signal with a 40 Hz low-pass Kaiser-windowed FIR filter (5 Hz transition); (3) calculated the mean and SD of ripple-band amplitude across all runs per channel. Candidate ripples were defined as periods when the squared amplitude exceeded 4 SD above the mean, with start/end points marked by crossings of 2 SD. Events lasting < 20 ms or > 200 ms were excluded. Ripple peaks were identified with MATLAB’s findpeaks.m, and events with < 30 ms peak-to-peak intervals were merged.

To validate the results from the above ripple detection approach, we performed a validity check on the detected ripples using the unfiltered raw signal. For each detected ripple, we identified the local voltage peaks within the event duration in the raw data and calculated the corresponding Inter-Peak Intervals (IPIs). True ripple events should be manifested as multi-peak oscillatory waveforms in the raw trace, whose IPIs should roughly correspond to the targeted ripple frequency range. In addition, we have performed an alternative ripple detection method following previous research (Vaz et al., 2019). Specifically, we first bandpass-filtered the iEEG signal in the ripple range (80-120 Hz) using a second-order Butterworth filter and then applied a Hilbert transform to extract the instantaneous amplitude. Candidate events were identified when the Hilbert envelope exceeded 2 SD above the mean amplitude. Events were kept as ripples only if they lasted at least 25 ms and reached a peak amplitude >3 SD. Adjacent events separated by <15 ms were merged.

To prevent contamination from pathological activity, interictal epileptiform discharges (IEDs) were rigorously screened. Channels within clinically identified epileptogenic zones were excluded. For the remaining channels, IEDs were independently visually inspected, marked, and verified by two neurologists at Beijing Xuanwu Hospital before the ripple detection analysis. In addition, IEDs typically exhibit broadband power increases (1-180 Hz), whereas physiological ripples show narrow high-frequency activity (70-180 Hz) and were further confirmed by performing the time-frequency analysis in Brainstorm (https://neuroimage.usc.edu/brainstorm/). Candidate ripple events occurring within 50 ms of an IED duration range were excluded.

To obtain the peri-ripple spectrograms, we first epoched iEEG data into 6-s segments, ranging from 3 s before the ripple peak to 3 s after it. The epoched data were then convolved with complex Morlet wavelets (six cycles) in the range of 1 to 250 Hz, with 2 Hz steps. The resulting complex wavelet transform was squared to obtain spectral power. Power values were z-transformed for each frequency and channel using the mean and the SD of the power across all epochs belonging to the same experimental run. Final spectrograms were centered around ripple peaks (± 250 ms) in 500-ms windows. All iEEG analyses were performed in MATLAB using custom scripts and functions from the FieldTrip toolbox (Oostenveld et al., 2011).

### Analysis of ripple rate changes and cluster-based permutation test

Detected ripples were time-locked to stimulus onset for the encoding and maintenance stages, and to behavioral responses for the retrieval stage. For each trial, ripple rate was calculated by dividing the number of ripples by the duration (in seconds) of each task stage: encoding (0-3 s post-stimulus onset), maintenance (3-10 s), and retrieval (from probe onset to response). The ripple rate in a pre-encoding time window (200 to 800 ms before stimulus onset) was also computed and served as the baseline.

We used linear mixed-effects model (LMM) analysis to compare the ripple rates between each VSTM task stage and the pre-encoding baseline, as well as between different task stages. The analysis was performed using the R package lme4 (Brown, 2021). The models were specified as follows:

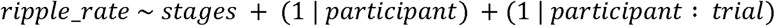

Here, ripple_rate refers to the ripple rate per channel, and stages is a fixed effect with four levels: pre-encoding baseline, encoding, maintenance, and retrieval. Random intercepts were included for each participant and for trials nested within participants to account for within-subject and within-trial variability. We then conducted post hoc pairwise contrasts between estimated marginal means (emmeans) of the four stages to test whether ripple rates differed between specific stages. The resulting *p*-values from these pairwise contrast tests were corrected for multiple comparisons using the Benjamini-Hochberg false discovery rate (FDR) procedure (Benjamini and Hochberg, 1995).

To quantify ripple rate changes at a finer temporal resolution, we computed ripple rates using non-overlapping 100-ms time windows within each task stage. The resulting time series was smoothed using a 400-ms Gaussian kernel. Ripple rates were then compared to the pre-encoding baseline separately for the encoding, maintenance, and retrieval stages. To correct for multiple comparisons, we performed non-parametric cluster-based permutation tests using the MATLAB code (https://doi.org/10.5281/zenodo.10877825). Specifically, for each time point, we computed the empirical ripple rate difference between conditions (e.g., encoding vs. baseline). We then compared it to a null distribution of ripple rate differences, which were generated by randomly permuting condition labels (e.g., encoding vs. baseline) 5,000 times. At each iteration, the ripple rate difference between conditions was recalculated. The empirical threshold for significance (α = 0.05, two-tailed) was determined from this null distribution. Time points of empirical ripple rate difference exceeding this threshold were grouped into clusters. For each cluster, a cluster-level statistic was computed by summing the ripple rate differences within the cluster. The observed cluster statistics were then compared against the distribution of maximum and minimum cluster-level statistics derived from the 5,000 permutations to determine significance. This same cluster-based permutation procedure was also used to compare ripple rates between remembered and forgotten trials, and between fast and slow remembered trials.

### Ramping-up effects analysis

To test whether the ramping-up of ripple activity during maintenance predicted VSTM performance, we fit generalized linear mixed-effects models (GLMMs) with a Poisson link function. The main model was specified as:

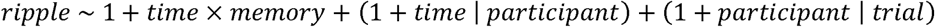

Here, *ripple* refers to the trial-level ripple count (Poisson-distributed). *Time* corresponds to the 7-s maintenance time index, and *memory* reflects VSTM accuracy (remembered = 1, forgotten = 0) or response speed (fast vs. slow), entered as fixed factors. Random intercepts and time slopes were included for each participant, and random intercepts for trials nested within participants. When this time × memory interaction effect was significant, we conducted follow-up analyses by fitting separate GLMMs for remembered and forgotten trials (or fast and slow trials). These post-hoc models tested whether ripple activity increased over time within each memory outcome, and corresponding *p*-values were further FDR corrected.

To validate our GLMM results, we also performed a group-level ramping-up analysis. Ripple rates were first averaged across trials and channels, yielding a time series of mean ripple rates during the maintenance stage for each participant. A linear regression was then applied to each participant’s ripple rate time series, with the regression slope quantifying the change in ripple rates over time. Independent *t*-tests were used to assess whether slopes were significantly greater than zero for remembered and forgotten trials, and to compare slopes between these two conditions.

In addition, to test whether the ripple dynamics differ in predicting VSTM and LTM performance, we fitted a GLMM, with a three-way time × memory × test_type interaction, in which the test_type refers to VSTM and LTM tests:

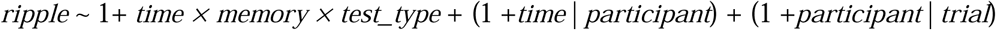

To further test whether the ripple dynamics differ between hippocampus and LTL, we fitted a GLMM, with a three-way time × region × memory interaction, in which region refers to hippocampus and LTL:

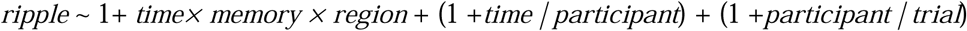

### Coupled ripple analysis

We first examined the temporal coordination between HPC and LTL ripple rates in a broader time window (i.e., [-0.5 s, 0.5 s]), which was employed to examine the lead-lag between the ripples of two regions. First, the hippocampus channels were paired with all LTL channels in each participant. For each HPC-LTL channel pair, we computed the hippocampal ripple peak-locked LTL ripple rate in this broader time window centered on Hippocampal (HPC) ripple peaks (Dickey et al., 2022). Additionally, non-ripple surrogates were derived for each channel pair, and the surrogate-locked LTL ripple rates were computed in time windows of the same length. Specifically, for all the *n* hippocampal ripples, we randomly selected *n* time points in the hippocampal channel with no ripple occurring 0.5 s before and after. Then, the LTL ripples were aligned to these non-ripple surrogates to obtain the surrogate-locked LTL ripple rates. This procedure was repeated 1000 times to generate a surrogate distribution, and empirical hippocampal ripple-locked LTL rates were z-scored relative to it, resulting in normalized LTL ripple rates. The normalized LTL ripple rates were then averaged across all HPC-LTL channel pairs for each participant. Cluster-based permutation tests assessed whether normalized LTL ripple rates exceeded zero within the ± 0.25 s window across participants. This analysis was conducted across all task stages and within each of the task stages (i.e., encoding, maintenance, and retrieval).

Next, for the coupled ripples, we identified LTL ripples that temporally overlapped with or occurred within ± 50 ms of a hippocampal ripple peak for each HPC-LTL channel pair following previous research (Vaz et al., 2019). The coupled ripple counts were then pooled across all trials and channel pairs within each participant, and subsequently averaged across participants. On average, each participant exhibited 439 (SD:125) coupled ripples in the VSTM task. When broken down by stage, each participant showed 143 ± 45 coupled ripples during encoding, 274 ± 76 during maintenance, and 23 ± 6 during retrieval. Coupled ripple rates were then computed and contrasted between remembered and forgotten trials across the task and during each task stage.

To rule out the possibility that the observed difference in coupled ripple rates between remembered and forgotten trials was driven by unequal trial numbers (i.e., a greater number of remembered trials), we performed a bootstrapping analysis. Specifically, for each participant, we randomly resampled remembered trials with replacement 1,000 times, matching the number of remembered trials to the number of forgotten trials (see Tables S2). For each bootstrap sample, we computed coupled ripple rates separately for remembered and forgotten trials. Statistical significance was assessed based on the proportion of bootstrap samples in which the coupled ripple rate was higher in remembered than in forgotten trials.

### Multi-variate decoding of VSTM representations

To decode category-specific memory representations, we trained a linear support vector machine (SVM) to classify the four picture categories (animal, fruit, electrical device, furniture) from remembered and forgotten trials. The chance level for decoding was set at 25%. Inspired by prior studies showing that memory content is best captured by a broad range of spectral power (Staresina et al., 2016; Yaffe et al., 2014), we extracted broadband (2-180 Hz) spectral power from hippocampal and LTL channels. Time-frequency transformation was performed using complex Morlet wavelets (2-29 Hz in 1-Hz steps and 30-180 Hz in 5-Hz steps, 6 cycles), and all power spectral data were down-sampled to 100 Hz after time-frequency transformation. Spectral power was z-scored for each frequency and channel across runs. To obtain time-resolved memory reactivation, decoding was performed in sliding time windows for both the training and test data, with a 400-ms window length and a 50-ms increment.

To increase the signal-to-noise ratio, the spectral power was averaged across time points within each time window, resulting in frequency × channel features as input to the SVM model. For each participant, normalized power from all hippocampus or LTL channels was used as input to the SVM. A leave-one-trial-out cross-validation was used to estimate decoding accuracy. SVMs were trained separately on each time window during encoding (yielding 60 decoders) and tested across all time windows of the task to assess temporal generalization. Decoding accuracy at each time point was statistically compared against the 25% chance level using the cluster-based permutation test.

### Coupled ripple-locked decoding accuracy

To assess whether memory reinstatement was linked to coupled ripples between the hippocampus and LTL, we computed decoding accuracy time-locked to coupled ripple events. First, we trained category-level classifiers on broadband iEEG data at every single time window during the encoding stage (0-3 s) and tested them across all time points of both the encoding and maintenance phases using a leave-one-trial-out cross-validation method (see details in the multi-variate decoding of VSTM representations). Second, for every single trial, we extracted the continuous decoding accuracy time series locked to the peak of each coupled hippocampal ripple within a window of [-0.5 s, 0.5 s] following previous research (Zhang et al., 2018). Next, to determine if memory reactivation was significantly locked to these coupled ripple events, in addition to using the chance level (0.25) as a baseline, we compared this empirical ripple-locked decoding accuracy against a surrogate baseline (constructed by randomly shuffling non-coupled time points locked decoding accuracy for 1,000 times to create a z-score distribution). A cluster-based permutation test was then performed across the [-0.5 s, 0.5 s] time window. The “identified cluster” is the resulting significant time window around the ripple peak where decoding accuracy was significantly elevated above the surrogate baseline.

As a control, we examined decoding accuracy locked to independent hippocampal or LTL ripples (i.e., ripples not overlapping or occurring within 50 ms of a ripple in the other region). Decoding accuracy for these independent ripples was z-scored relative to a non-ripple surrogate distribution, and the same statistical procedures were applied.

## Acknowledgement

We thank the patients who volunteered to participate in the experiments and the support from our collaborators from Beijing Xuanwu Hospital, who provided the patient resources. This work was supported by the STI 2030-Major Projects (2021ZD0200401, 2021ZD0200409), the Fundamental Research Funds for the Central Universities (226-2024-00118), National Natural Science Foundation of China (32100851), and Guangdong Basic and Applied Basic Research Foundation (No. 2024A1515012667 and No. 2023A1515110311). This work was also supported by the grant from the Research Center for Brain Cognition and Human Development, Guangdong, China (No. 2024B0303390003).

## Author contributions

J.L. and G.X. conceived the original experiment. J.L., X.H., C.Y., N.A., S.Z., and Y.C. performed the analysis. J.L., X.H., and Y.C. wrote the initial manuscript. J.L., X.H., C.Y., N.A., G.X., S.Z., and Y.C. revised the manuscript.

## Data availability

All preprocessed data and custom MATLAB code necessary to reproduce the main conclusions of this study are available on the Open Science Framework: https://osf.io/gwe62/?view_only=39b3ddbeaae342c8a2696a1e16473712.

Any additional information required to reanalyze the data reported in this paper is available from the lead contact upon request.

