## Supplemental Material for "Ramping-up hippocampal ripples and their neocortical coupling support human visual short-term memory"

This file contains Figures S1-S10 and Tables S1-S2.

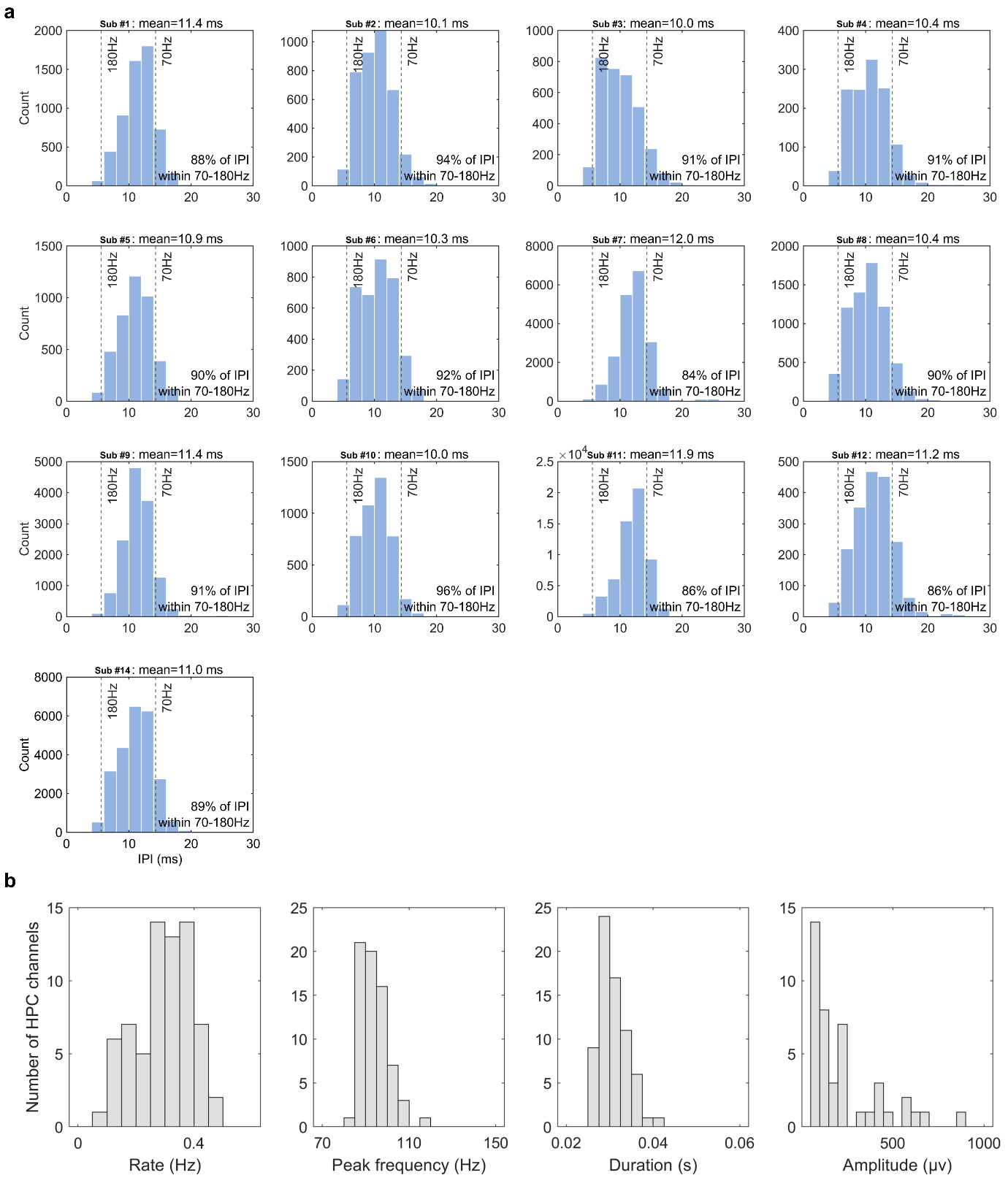

**Figure S1. Properties of ripples in HPC channels. (a)** Inter-peak interval (IPI) distributions of unfiltered raw signals within detected hippocampal ripple epochs. Individual histograms represent data from each of the 13 participants. **(b)** Distributions of ripple rate, peak frequency, duration, and amplitude across all HPC channels.

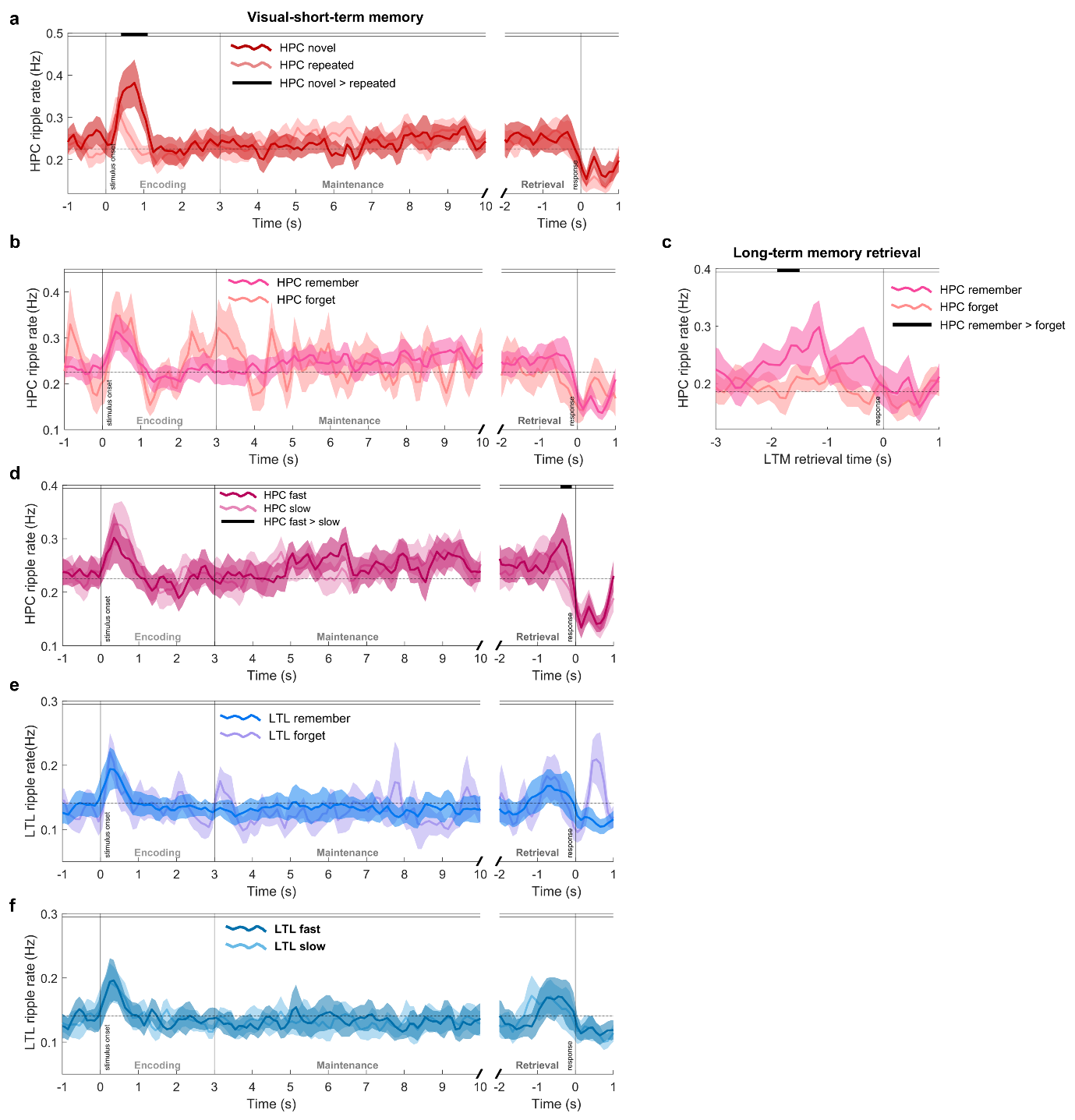

**Figure S2. Ripple rates in the hippocampus and lateral temporal lobe (LTL) during the visual short-term memory (VSTM) task and long-term memory (LTM) retrieval. (a)** Time course of hippocampal ripple rates for novel (first presentation) and repeated (second and third presentations) trials. The black horizontal bar indicates a significant cluster (500–1100 ms post-stimulus) with higher ripple rates for novel versus repeated trials (*p*_cluster_ < 0.001). **(b)** No significant differences were observed between hippocampal ripple rates for remembered and forgotten trials in any VSTM stage (all *p*s_cluster_ > 0.466). **(c)** Hippocampal ripple rates were significantly higher for LTM remembered than forgotten trials within a cluster (i.e., 1600-1900 ms before LTM retrieval responses) (*p*_cluster_ = 0.047). **(d)** Hippocampal ripple rates were significantly higher for fast compared with slow trials within a pre-response cluster (i.e., 150-450 ms before retrieval responses, *p*_cluster_ = 0.014). **(e)** No significant differences were observed between LTL ripple rates of remembered versus forgotten trials in any VSTM stage (all *p*s_cluster_ > 0.274). **(f)** No significant differences were observed between hippocampal ripple rates for fast and slow remembered trials in any VSTM stage (all *p*s_cluster_ > 0.237). Shaded areas represent ± 1 SEM.

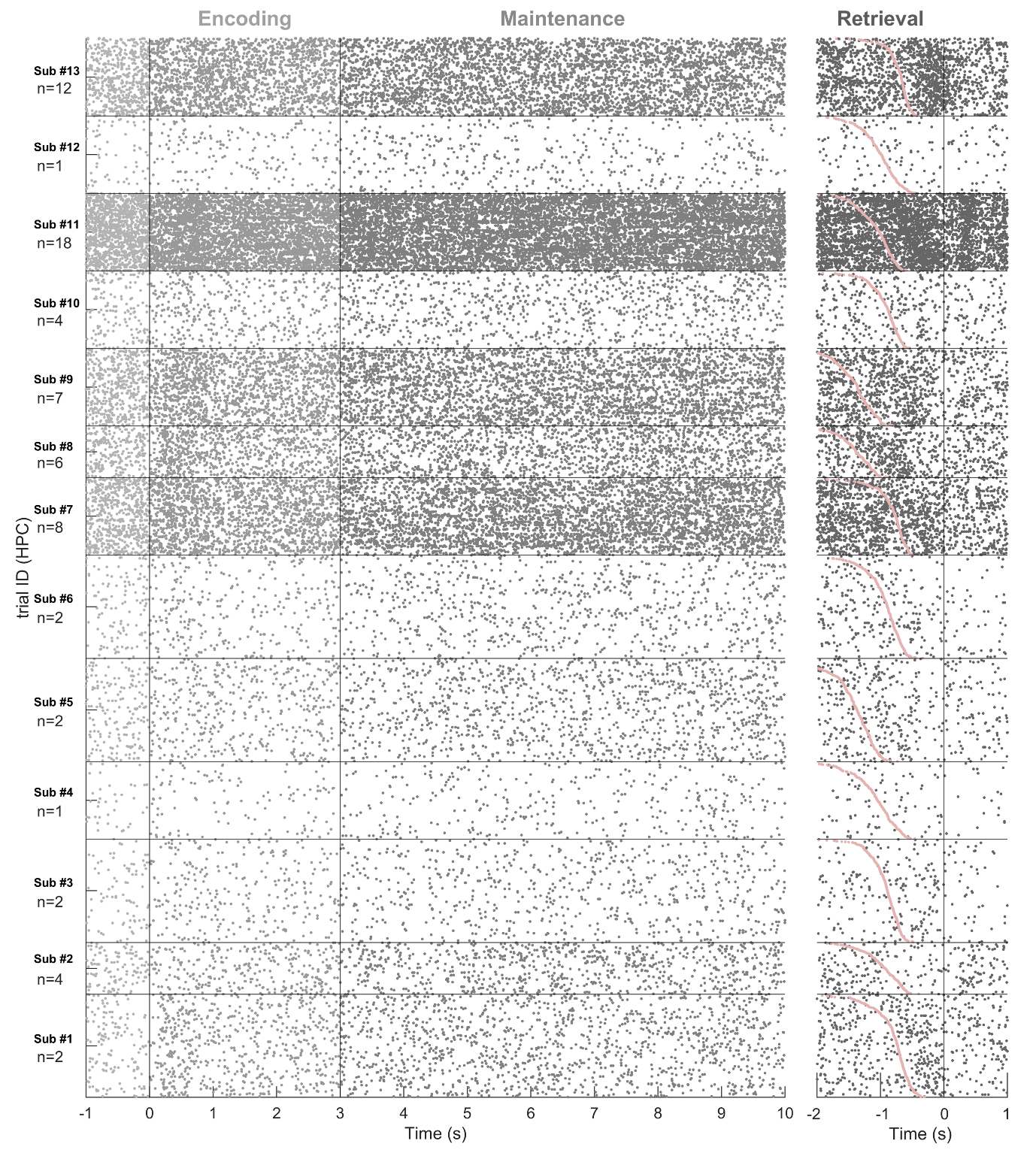

**Figure S3. Raster plot of hippocampal ripples for individual participants.** Each row represents a single VSTM trial, sorted by reaction time, with each dot marking the peak time of a ripple. For participants with multiple hippocampal channels, ripple events detected across all channels are overlaid within the same trial row; thus, participants with more hippocampal channels show a higher spatial density of plotted dots. The three vertical lines from left to right indicate onsets of encoding, maintenance, and retrieval response, respectively. The bolded pink curve represents the probe onset of each trial during retrieval.

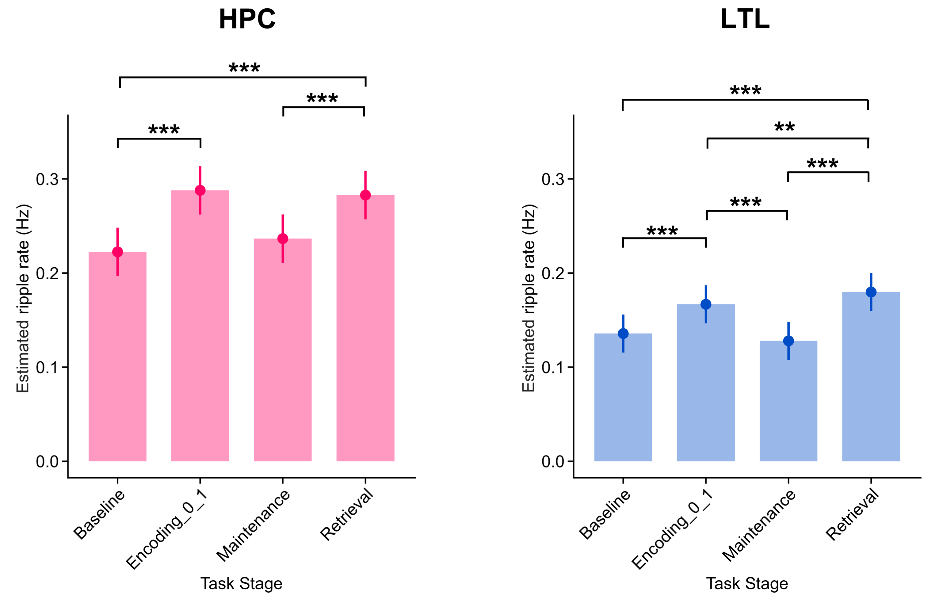

**Figure S4. Ripple rate comparison between task stages using a tuned encoding window (0-1 s post-stimulus onset), while keeping time windows of other stages unchanged.** Stage-averaged ripple rates estimated using Linear Mixed-Effects Models (LMMs) for hippocampal channels (left) and lateral temporal lobe (LTL) channels (right). *: *p*_FDR_ < 0.05, **: *p*_FDR_ < 0.01, ***: *p*_FDR_ < 0.001.

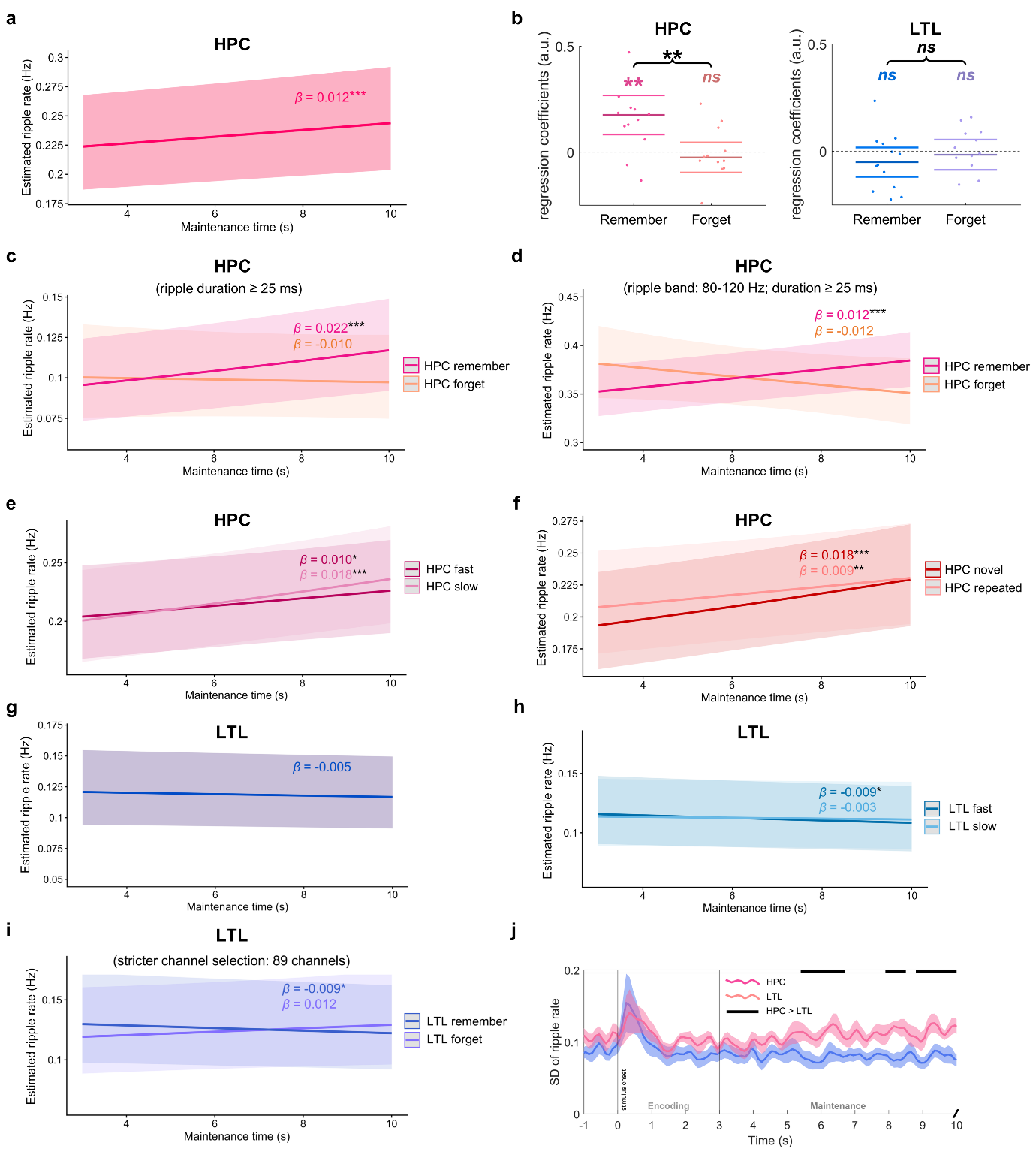

**Figure S5. Control analyses for the ripple ramping-up effects in the hippocampus (HPC) and lateral temporal lobe (LTL).** **(a)** Significant ramping-up of hippocampal ripple rates during the maintenance period across all trials, channels, and participants (*β* = 0.012, *z* = 4.678, *p* < 0.001). The y-axis shows model-estimated ripple rates. **(b)** Group-level analysis of the ripple ramping-up effects in the hippocampus and lateral temporal lobe (LTL). Left: for the hippocampus, remembered trials showed significantly positive slopes (*t*(12) = 3.580, *p* = 0.004), whereas forgotten trials did not (*t*(12) = -0.669, *p* = 0.516). The slopes for remembered trials were significantly greater than those for forgotten trials (*t*(12) = 3.400, *p* = 0.005). Right: for the LTL, neither remembered trials (*t*(12) = -1.408, *p* = 0.185) nor forgotten trials (*t*(12) = -0.437, *p* = 0.670) of the visual short-term memory (VSTM) task showed significant slopes against zero. The slopes did not differ between VSTM remembered and forgotten trials (*t*(12) = -0.707, *p* = 0.493). **(c)** Control analyses by restricting hippocampal ripples (as detected in the main text) to those with a duration ≥ 25 ms, the results found a significant time × VSTM accuracy interaction (*β* = 0.033, *z* = 2.983, *p* = 0.003). Further analyses revealed a significant ramping-up of ripple rates over the maintenance period for remembered trials (*β* = 0.022, *z* = 6.178, *p*_FDR_ < 0.001), but not for the forgotten items (*β* = -0.010, *z* = -0.874, *p*_FDR_ = 0.382), consistent with Fig. 3a in the main text. **(d)** Control analyses for hippocampal ramping-up effect using an alternative frequency range (80-120 Hz) and detection criteria (duration ≥ 25 ms and peak amplitude > 3 SD above baseline) following Vaz et al. (2019, *Science*). Our results found a significant time × VSTM accuracy interaction was again observed (*β* = 0.024, *z* = 3.281, *p* = 0.001). Further analyses revealed a significant ramping-up of ripple rates over the maintenance period for remembered trials (*β* = 0.012, *z* = 4.998, *p*_FDR_ < 0.001), but not for the forgotten items (*β* = -0.012, *z* = -1.648, *p*_FDR_ = 0.099), consistent with Fig. 3a in the main text. **(e)** Both fast and slow remembered trials showed significant hippocampal ramping-up effects (fast: *β* = 0.010, *z* = 2.463, *p*_FDR_ = 0.014; slow: *β* = 0.018, *z* = 4.625, *p*_FDR_ < 0.001), with no significant time × retrieval speed interaction (*β* = 0.009, *z* = 1.570, *p* = 0.117). The y-axis shows model-estimated ripple rates. **(f)** Both novel and repeated trials showed significant hippocampal ramping-up effects (novel: *β* = 0.018, *z* = 4.00, *p*_FDR_ < 0.001; repeated: *β* = 0.009, *z* = 2.973, *p*_FDR_ = 0.003), with no significant time × novelty (novel vs. repeated) interaction effect (*β* = -0.009, *z* = -1.716, *p* = 0.086). **(g)** Across all trials, channels, and participants, a linear mixed-effects model revealed a non-significant trend of decreasing LTL ripple rates during the maintenance period (*β* = -0.005, *z* = -1.772, *p* = 0.076). The y-axis shows model-estimated ripple rates. **(h)** LTL ripple rates decreased significantly for the fast trials, with a similar but non-significant trend for the slow trials (fast: *β* = -0.009, *z* = -2.245, *p*_FDR_ = 0.049; slow: *β* = -0.003, *z* = -0.721, *p*_FDR_ = 0.471) and no significant time × retrieval speed interaction (*β* = 0.006, z = 1.087, *p* = 0.277). The y-axis shows model-estimated ripple rates. **(i)** Control analyses when restricting analyses to LTL bipolar channel pairs with at least one contact located within LTL gray matter or within 2 mm of gray matter (n = 89 channels). The results revealed a significant time × VSTM accuracy interaction (*β* = -0.020, *z* = -2.025, *p* = 0.043). Further analyses showed that this interaction was driven by a significant decrease in ripple rates for remembered items (*β* = -0.009, *z* = -2.641, *p*_FDR_ = 0.017), whereas no significant effect was observed for forgotten trials (*β* = 0.012, *z* = 1.134, *p*_FDR_ = 0.257). **(j)** The standard deviation of hippocampal ripple rates was significantly greater than that of LTL channels during the maintenance and pre-retrieval response periods across participants (all *p*s_cluster_ < 0.032), suggesting a greater heterogeneity for hippocampal channels than LTL channels. After controlling for the standard deviation for time and regions, the interaction of *Time* and *Region* in predicting $Ripple\_rate$ remained significant (*p* = 0.017), indicating that the observed difference in ramping-up effects between HPC and LTL cannot be simply attributed to regional heterogeneity. Significant clusters are indicated by horizontal black bars. *: *p*_FDR_ < 0.05, *ns*: not significant. *: *p* (*p*_FDR_) < 0.05, **: *p* (*p*_FDR_) < 0.01, ***: *p* (*p*_FDR_) < 0.001.

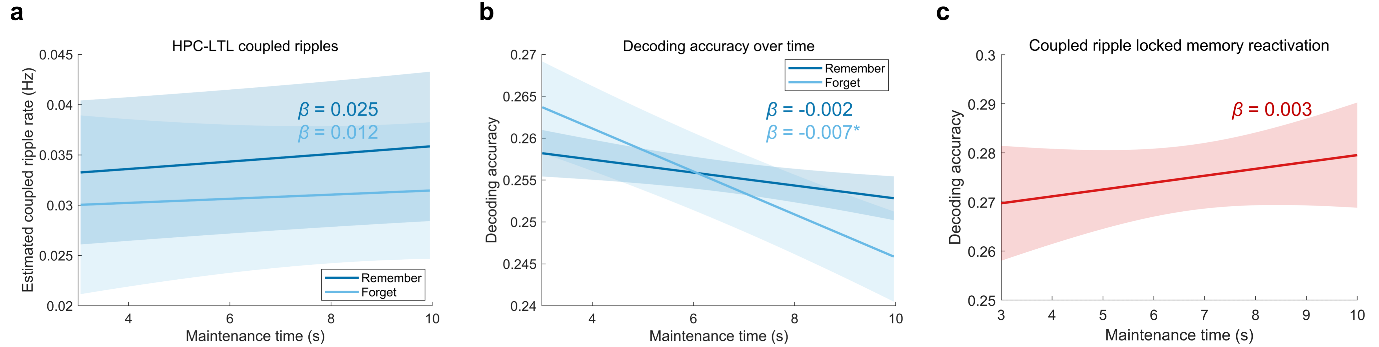

**Figure S6. Temporal dynamics of coupled ripples and decoding accuracy across the maintenance interval.** **(a)** Coupled ripple rates over the 7-second maintenance period for VSTM remembered and forgotten trials. Coupled ripples showed a non-significant ramping-up pattern for both remembered and forgotten trials (*p*s_FDR_ > 0.281), and there was no significant time x memory interaction (*p* = 0.506, see Fig. S6 below). **(b)** Decoding accuracy over the maintenance period, evaluated using the classifier trained on the late encoding window (1.85-2.25 s). The results revealed a significant time * memory accuracy interaction (*p* = 0.009, see Fig. S6 below). Further analysis revealed that this interaction was driven by a significant decline in decoding accuracy on forgotten trials (*p*_FDR_ = 0.031), whereas decoding accuracy on remembered trials was maintained (*p*_FDR_ = 0.214). **(c)** Coupled ripple-locked decoding accuracy over the maintenance period. The results showed a non-significant numeric increase across the maintenance period (*p* = 0.586). Error bands represent ± 1 SEM. *: *p*_FDR_: < 0.05.

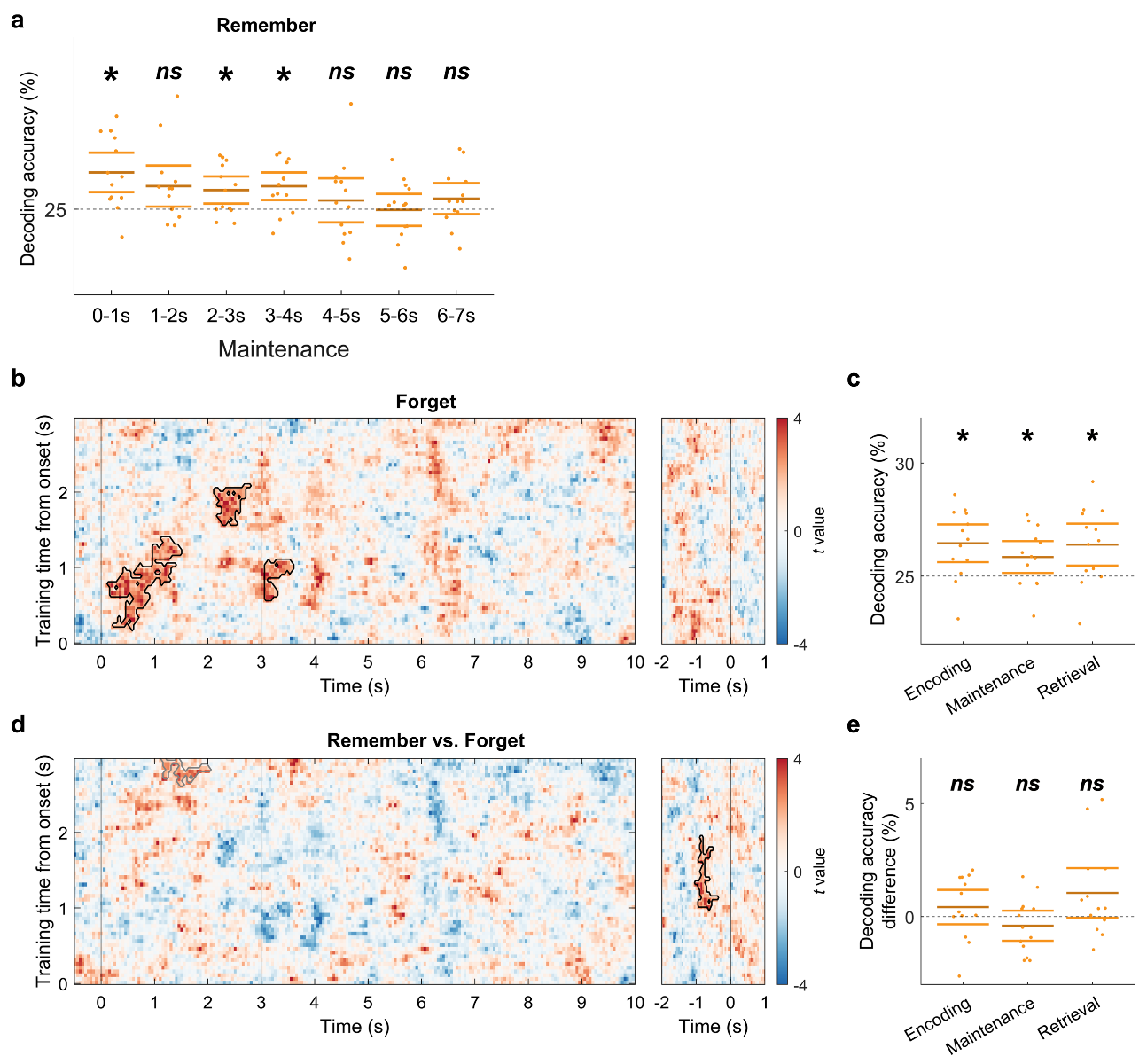

**Figure S7. Control analyses for decoding accuracy in the lateral temporal lobe (LTL) for remembered and forgotten trials. (a)** Decoding accuracy was computed separately for each 1-s time bin of the maintenance period for remembered trials, with the first, third, and fourth time bins significant above chance (i.e., 25%) (*p*s_FDR_ < 0.042). **(b)** Decoding accuracy of forgotten trials compared to chance level (25%) across the task (left: encoding and maintenance. Clusters showing significantly above-chance decoding accuracy are circled by black lines (all *p*s_cluster_ < 0.028). **(c)** Mean decoding accuracy across all time windows within each stage (encoding, maintenance, retrieval) was significantly above chance across participants (all *p*s_FDR_ < 0.037). **(d)** Decoding accuracy for VSTM remembered trials was significantly higher than for forgotten trials during retrieval in the cluster circled by black lines (*p*_cluster_ = 0.012), with a similar trend but non-significant cluster during encoding (*p*_cluster_ = 0.064, circled by grey lines). **(e)** Mean decoding accuracy across all time windows within each stage did not differ significantly between VSTM remembered and forgotten trials (all *p*s_FDR_ > 0.258). *: *p*_FDR_ < 0.05, *ns*: not significant.

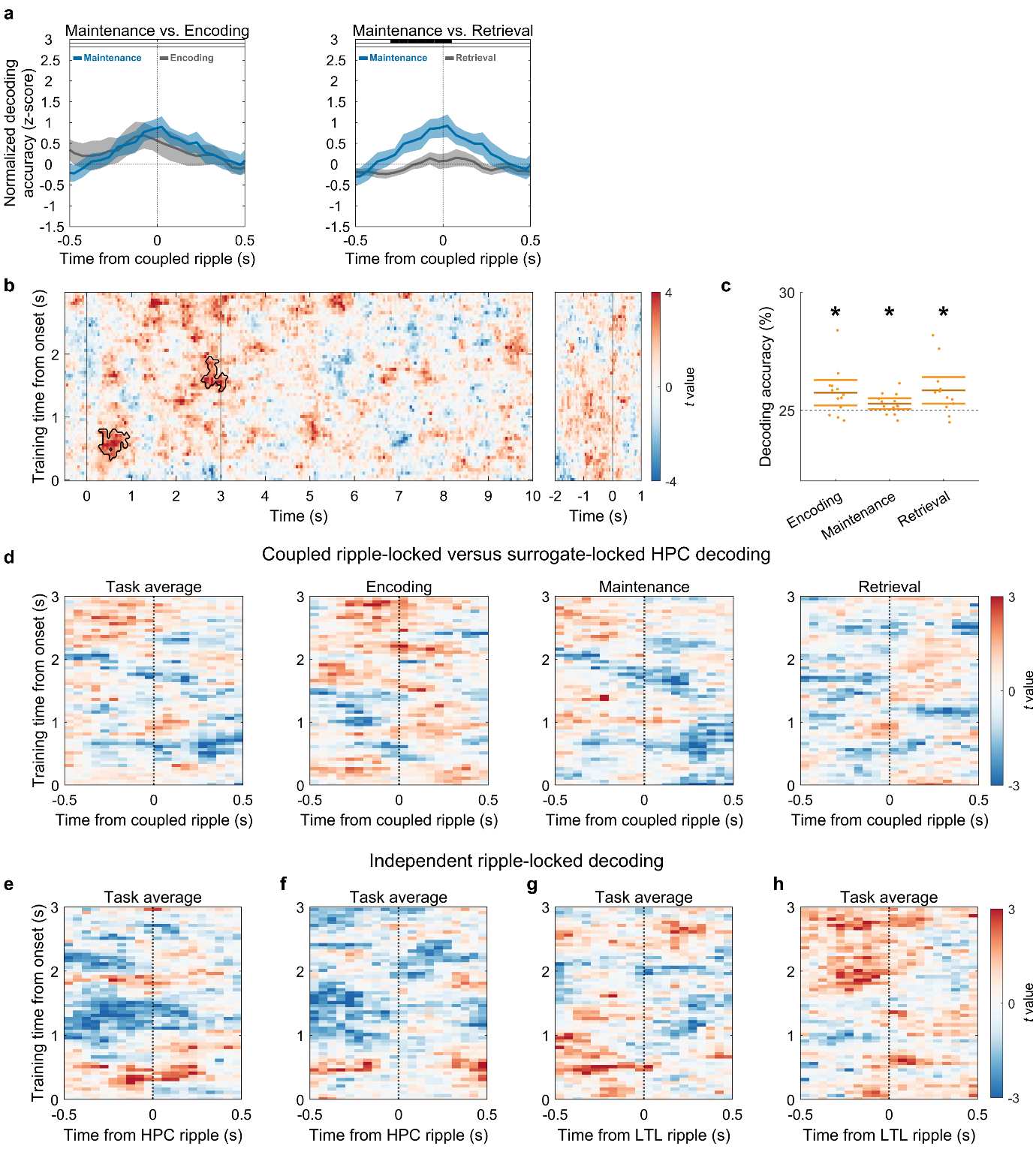

**Figure S8. Control analyses for coupled ripple-locked memory reactivation. (a)** Contrasting the coupled ripple-locked lateral temporal lobe (LTL) decoding accuracy during maintenance with that during encoding and retrieval, respectively. The black horizontal bar indicates a significant cluster with decoding accuracy for maintenance compared to the other task stage (*p*_cluster_ < 0.05). **(b)** Hippocampal (HPC) decoding accuracy compared to chance level (25%). Clusters indicate time windows with significantly above-chance decoding (*p*s_cluster_ < 0.031) and are circled by black lines. **(c)** Averaged HPC decoding accuracies across all train-test time windows within individual task stages were all significantly above chance (*p*s_FDR_ < 0.038). **(d)** No significant clusters were found for HPC decoding accuracy locked to HPC-LTL coupled ripples across all task stages or within any task stage (all *p*s_cluster_ > 0.470). **(e-f)** Neither HPC (e) nor LTL (f) decoding accuracy was significantly locked to independent HPC ripples (all *p*s_cluster_ > 0.222). **(g-h)** Neither HPC (g) nor LTL (h) decoding accuracy was significantly locked to independent LTL ripples (all *p*s_cluster_ > 0.203). *: *p*_FDR_ < 0.05.

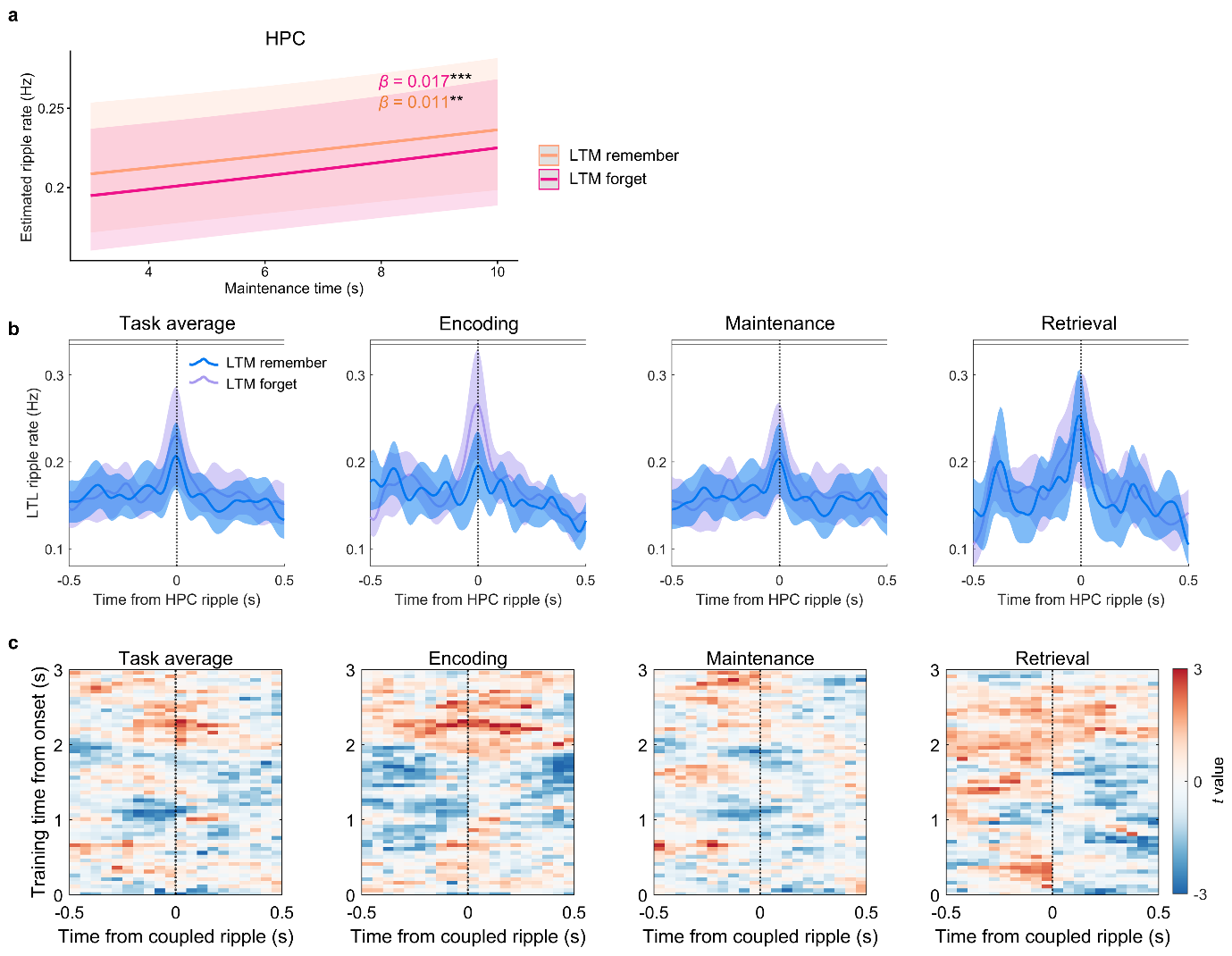

**Figure S9. Control analyses for ruling out long-term memory (LTM) confounds in hippocampal ramping-up effects, coupled ripples, and ripple-locked memory reactivation.** **(a)** Hippocampal (HPC) ripples showed significant ramping-up effects for both subsequently LTM-remembered (*β* = 0.017, *z* = 4.141, *p*_FDR_ < 0.001) and LTM-forgotten (*β* = 0.011, *z* = 2.987, *p*_FDR_ = 0.003) trials, with no significant time × LTM accuracy interaction (*β* = 0.003, *z* = 0.439, *p* = 0.661). **(b)** HPC-LTL coupled ripple rates did not differ between subsequently LTM remembered and forgotten trials (*p*s_cluster_ > 0.626). **(c)** Coupled ripple-locked memory reactivation also showed no significant difference between subsequently LTM remembered and forgotten trials (all *p*s_cluster_ > 0.135). **: *p*_FDR_ < 0.01, ***: *p*_FDR_ < 0.001.

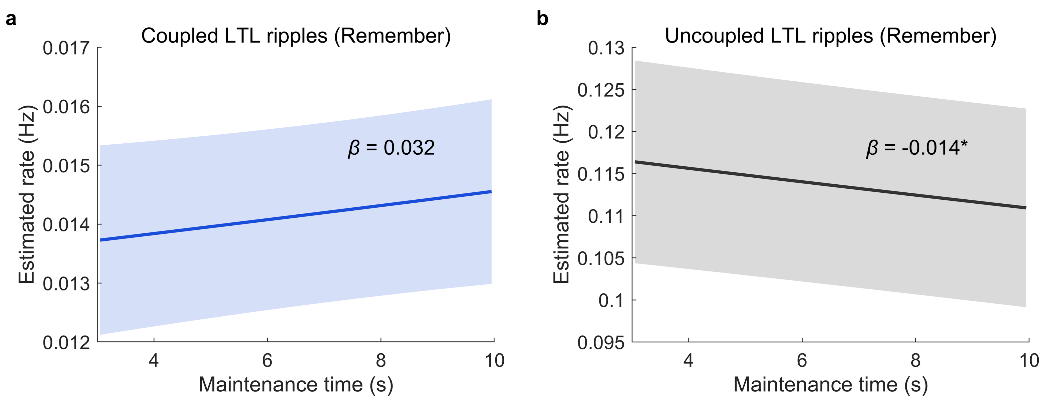

**Figure S10. Temporal dynamics of lateral temporal lobe (LTL) ripples that were coupled (a) or uncoupled to the hippocampal (HPC) ripples (b).** The results revealed a non-significant, positive numerical trend for LTL ripples that coupled to HPC ripples (*β* = 0.032, *p* = 0.126), whereas uncoupled LTL ripples showed a significant decrease over the maintenance period (*β* = -0.014, *p* = 0.026), with the interaction effect showing a non-significant trend of greater ramping-up for coupled versus uncoupled LTL ripples (*β* = 0.031, *p* = 0.088). *: *p* < 0.05.

**Table S1. Participant-level summary of regional channel counts and ramping-up coefficients (*β*) for remembered and forgotten trials.** HPC = Hippocampus; LTL = Lateral Temporal Lobe; AMY = Amygdala; PHC = Parahippocampal Cortex; ERC = Entorhinal Cortex.

| Participant ID | HPC  (N_sub_ = 13) | LTL  (N_sub_ = 13) | AMY  (N_sub_ = 9) | PHC  (N_sub_ = 5) | ERC  (N_sub_ = 1) | HPC: *β*  (remember) | HPC: *β*  (forget) |
| --- | --- | --- | --- | --- | --- | --- | --- |
| Sub1 | 2 | 22 | 1 | 1 | / | -0.134 | 0.005 |
| Sub2 | 4 | 16 | / | 2 | / | 0.129 | -0.047 |
| Sub3 | 2 | 22 | 2 | 1 | / | 0.179 | -0.017 |
| Sub4 | 1 | 7 | / | / | / | 0.152 | -0.041 |
| Sub5 | 2 | 3 | 2 | / | 2 | 0.471 | 0.147 |
| Sub6 | 2 | 2 | / | / | / | 0.209 | 0.115 |
| Sub7 | 8 | 2 | 1 | / | / | 0.503 | -0.257 |
| Sub8 | 6 | 21 | 1 | / | / | 0.201 | 0.228 |
| Sub9 | 7 | 7 | 5 | 1 | / | 0.185 | -0.041 |
| Sub10 | 4 | 2 | 2 | / | / | 0.262 | -0.080 |
| Sub11 | 18 | 5 | 3 | / | / | 0.122 | -0.075 |
| Sub12 | 1 | 12 | 3 | / | / | -0.061 | -0.240 |
| Sub13 | 12 | 11 | / | 1 | / | 0.060 | -0.023 |
| N_ele | 69 | 132 | 20 | 6 | 2 | / | / |

**Table S2. Participant-level summary of trial number counts for remembered and forgotten trials.**

| Participant ID | Trial number  (Remember) | Trial number  (Forget) |
| --- | --- | --- |
| Sub1 | 309 | 27 |
| Sub2 | 149 | 19 |
| Sub3 | 298 | 38 |
| Sub4 | 229 | 23 |
| Sub5 | 265 | 71 |
| Sub6 | 302 | 34 |
| Sub7 | 232 | 20 |
| Sub8 | 152 | 16 |
| Sub9 | 212 | 40 |
| Sub10 | 229 | 23 |
| Sub11 | 228 | 24 |
| Sub12 | 235 | 17 |
| Sub13 | 242 | 10 |
